# Embryo mechanics cartography: inference of 3D force atlases from fluorescence microscopy

**DOI:** 10.1101/2023.04.12.536641

**Authors:** Sacha Ichbiah, Fabrice Delbary, Alex McDougall, Rémi Dumollard, Hervé Turlier

**Affiliations:** Center for Interdisciplinary Research in Biology (CIRB), Collège de France, CNRS, INSERM, Université PSL, 11 place Marcelin Berthelot, Paris, France; Laboratoire de Biologie du Développement de Villefranche-sur-Mer (LBDV), Institut de la Mer de Villefranche, Sorbonne Université, CNRS, 181 chemin du Lazaret, Villefranche-sur-Mer, France

**Keywords:** force inference, inverse problem, foam, triangle mesh, surface tension, Laplace pressure, embryo, tissue mechanics

## Abstract

The morphogenesis of tissues and embryos results from a tight interplay between gene expression, biochemical signaling and mechanics. Although sequencing methods allow the generation of cell-resolved spatio-temporal maps of gene expression in developing tissues, creating similar maps of cell mechanics in 3D has remained a real challenge. Exploiting the foam-like geometry of cells in embryos, we propose a robust end-to-end computational method to infer spatiotemporal atlases of cellular forces from fluorescence microscopy images of cell membranes. Our method generates precise 3D meshes of cell geometry and successively predicts relative cell surface tensions and pressures in the tissue. We validate it with 3D foam simulations, study its noise sensitivity, and prove its biological relevance in mouse, ascidian and *C. elegans* embryos. 3D inference allows us to recover mechanical features identified previously, but also predicts new ones, unveiling potential new insights on the spatiotemporal regulation of cell mechanics in early embryos. Our code is freely available and paves the way for unraveling the unknown mechanochemical feedbacks that control embryo and tissue morphogenesis.

## 1 Introduction

Understanding the mechanical regulation of embryo and tissue shape emergence is a longstanding goal in developmental biology and biological physics. Although gene expression patterning in early embryos is increasingly documented thanks to recent single cell sequencing methods [1, 2], we still know very little about how cellular forces are spatio-temporally patterned within embryos and tissues. This is due to the lack of efficient methods for extracting cell- and time-resolved mechanics in a systematic, tissue-wide, and noninvasive manner.

Most experimental methods to measure mechanics are local and time-consuming, such as micropipette aspiration, AFM measurement, or embedded droplet deformation [3–15], making the generation of spatio-temporal maps of mechanics tedious; others are invasive, such as laser ablation, perturbing normal tissue development [16, 17]; or they probe mechanics only at the tissue level [18–21], ignoring mechanical heterogeneities within the multicellular structure. Interestingly, all methods require live 3D imaging to follow the deformation of cells, tissues, or embedded objects. Advances in fluorescence microscopy allow us to record the geometry of cells during the development of an embryo *in toto* from the zygote to a few hundreds of cells with a confocal microscope [22] and up to thousands of cells with a light sheet microscope [23, 24]. Attractive new microscopy techniques have emerged to try to quantify cellular mechanics directly, such as Brillouin microscopy [25, 26], or membrane tension probes [27–29], but such methods still lack cross-validations and remain difficult to link directly to mechanical models of tissues.

An alternative idea that emerged a decade ago is to infer the forces that dictate the shape of cells directly from their geometry by solving an inverse mechanical model [30]. These mechanical inference methods (also called *force* or *stress inference*) are based only on image analysis and do not require tissue perturbation: they have therefore a lower entry barrier than many other methods, as they do not require complex experimental setups. They have been shown to be efficient in inferring tensions (and pressure) in 2D cell monolayers [31] and can be scaled to hundreds or thousands of cells. For tissues and embryos, most inference methods are based on the hypothesis that cells adopt shapes and arrangements similar to bubbles in a foam, as pointed out by D’Arcy Thompson more than a century ago [32]. This analogy implies that the mechanics of cells is dominated by tensile stresses on their surface, which are generated by actomyosin contractility [33]. Because actomyosin contractility may be regulated differentially in distinct cells or at different interfaces (such as the cell-medium and cell-cell interfaces [5]), embryos and tissues may be seen as heterogeneous foams, where each cellular interface may adopt a different tension. Actual inference methods also assume generally a quasistatic mechanical equilibrium, where the viscous relaxation of tensions (dozens of seconds) is much faster than typical developmental timescales (dozens of minutes to hours). This foam-like mechanical equilibrium underpins two force balances, the Young-Dupré and Young-Laplace equations (Section 2), relating surface tensions with contact angles and cell pressures with interface curvatures. In the next, we will therefore refer to tension and pressure inference.

First versions of tension inference methods [34, 35] neglected Laplace’s law by assuming straight cell interfaces, as in traditional vertex models [36, 37]. In addition, they treated tensions and pressure as independent variables, which made the inverse problem generally underdetermined and relatively sensitive to noise. Alternatively, segmentation of cell membranes into 2D polygonal lines to explicitly measure their curvature [38, 39] allows successive determinations of tension and pressure and makes the set of equations generally overdetermined. In the particular case where the whole tissue can be imaged with its boundaries - as this is generally the case for early embryos - the problem turns out to be systematically overdetermined. However the generalization of this approach to three dimensions has not been convincing so far, since high-quality images and a robust segmentation pipeline are required [40, 41]. To avoid such issue, an elegant variational 2D approach was recently proposed in which cell junctions are fitted by circular arcs to find tensions and pressure [42], taking advantage of a mapping between a heterogeneous 2D foam and the tiling of the space into ”circular arc polygons”. This tiling falls actually within the class of Möbius diagrams [43, 44], whose mapping to 2D foams was already pointed out mathematically [45]. In 3D however, interfaces have mean constant curvatures but are generally not portions of sphere and may adopt saddle-node shapes, as remarked for homogeneous foams already [46]. In contrary to a recent assumption [47], the generalization to 3D of the variational scheme developed by [42] with Möbius diagrams is mathematically not correct.

To fill the gap, we propose *foambryo* [48], a robust end-to-end computational method for performing tension and pressure inference in three dimensions, starting directly from 3D fluorescence microscopy of cell membranes. Our pipeline follows the 2D approach of [38], where we decouple the inferences of tensions and pressures. It relies particularly on a novel and efficient surface mesh reconstruction method to precisely quantify cell geometry. Our inversion algorithm exploits furthermore junction lengths and interface areas as weights to infer tensions and pressures more robustly. Importantly, we performed a comprehensive benchmarking of our pipeline using 3D foam-like mechanical simulations and a systematic sensitivity analysis on various equivalent force balance formulas. Our inference pipeline yields convincing results on early embryos of mice, worms and ascidians by recovering known mechanical characteristics and predicting new ones. We provide an easy-to-install Python package and a comprehensive set of user-friendly 3D visualization tools.

## 2 Results

### Delaunay-watershed algorithm for multimaterial mesh generation

An essential first step is to extract the precise geometry of cells from microscopy images. Voxel-based segmentation masks are heavy data structures that are not well adapted to measure geometrical features such as contact angles or mean curvatures. Alternatively, triangle mesh representations of cell interfaces possess several advantages: they are sparse data structures that facilitate the retrieval of geometric quantities using a discrete differential formulas[50, 51]. They are easy to render graphically and form basic elements for computational modeling, such as vertex models [52, 53] or finite element methods [54]. The surface meshes of interest in our case are triangular, nonmanifold to account for tricellular junctions, and multimaterial to keep track of the identity of each enclosed cell or region (”material”), in the spirit of [55]. Although triangle meshes can be generated by discretizing voxel-based segmentation masks directly, using marching cube algorithms [56] or more recent methods [57], we found that previous algorithms introduced large errors in angle measurements in general.

Therefore, we developed a novel algorithm that robustly generates nonmanifold multimaterial surface meshes from cell segmentation masks^1^. The first step consists of computing a Euclidean distance transform map (EDT) [59] from the cell segmentation mask ^2^, which represents a smooth topographic map of cell (and image) boundaries (Fig. 2a). From the distance map, we sample points at the extrema of the elevation value using a max-pooling operator, which serves as control points to generate a Delaunay tessellation of the space (triangulation in 2D or tetrahedralization in 3D). A dual Voronoi diagram is then generated from the Delaunay tessellation and is represented as an edge-weighted graph *𝒢* = (*𝒩, ε 𝒲*,), where *𝒩* is the set of nodes, representing tetrahedra in the dual space (triangles in 2D), the set of edges between these nodes *𝒲* and their associated weights. These weights are defined here according to the average value of the integrated distance map measured along the corresponding triangle (or edge in 2D) in the dual space (Extended data Fig. 2a). Seeding each region using masks, we partition this graph using a watershed algorithm [62] that separates the nodes in the graph between the different cells and the external region^3^. Mapped back on the dual Delaunay space, this partition defines a unique surface (contour in 2D) mesh that accurately follows cell boundaries. Our *Delaunay-watershed* mesh generation algorithm works just as well in 2D as in 3D (Fig. 2a). Since the main purpose of this mesh generation algorithm is to extract precise geometrical features, we generated a set of 47 foam-like simulations of embryos with a number of cells varying from 2 to 11, which we translated into artificial confocal fluorescent images of size ∼ [250 × 250 ×250] [66] to compare the error generated for different geometrical measures of interest (contact angles, mean curvature, junction length, area, and volume) by our pipeline and state-of-the-art surface meshing techniques implemented in CGAL [57]. Our Delaunay-watershed algorithm [67] outperforms CGAL [57] for the retrieval of contact angles (Fig. 2b), and cell volumes or junction lengths (Extended data Fig. 2b), while its precision is comparable for the retrieval of interface areas and mean curvatures (Extended data Fig. 2b).

### Tension and pressure balance

Once the geometry of the cells can be calculated from the cell segmentation mesh (Extended data Fig.1 and Supplementary Note), we have to formulate the inverse mechanical problem to retrieve the relative force of the cells from their geometry. A quasi-static foam-like equilibrium underpins two stress balance equations within the tissue (Fig. 1d). The Young-Laplace equation

**Fig. 1:**
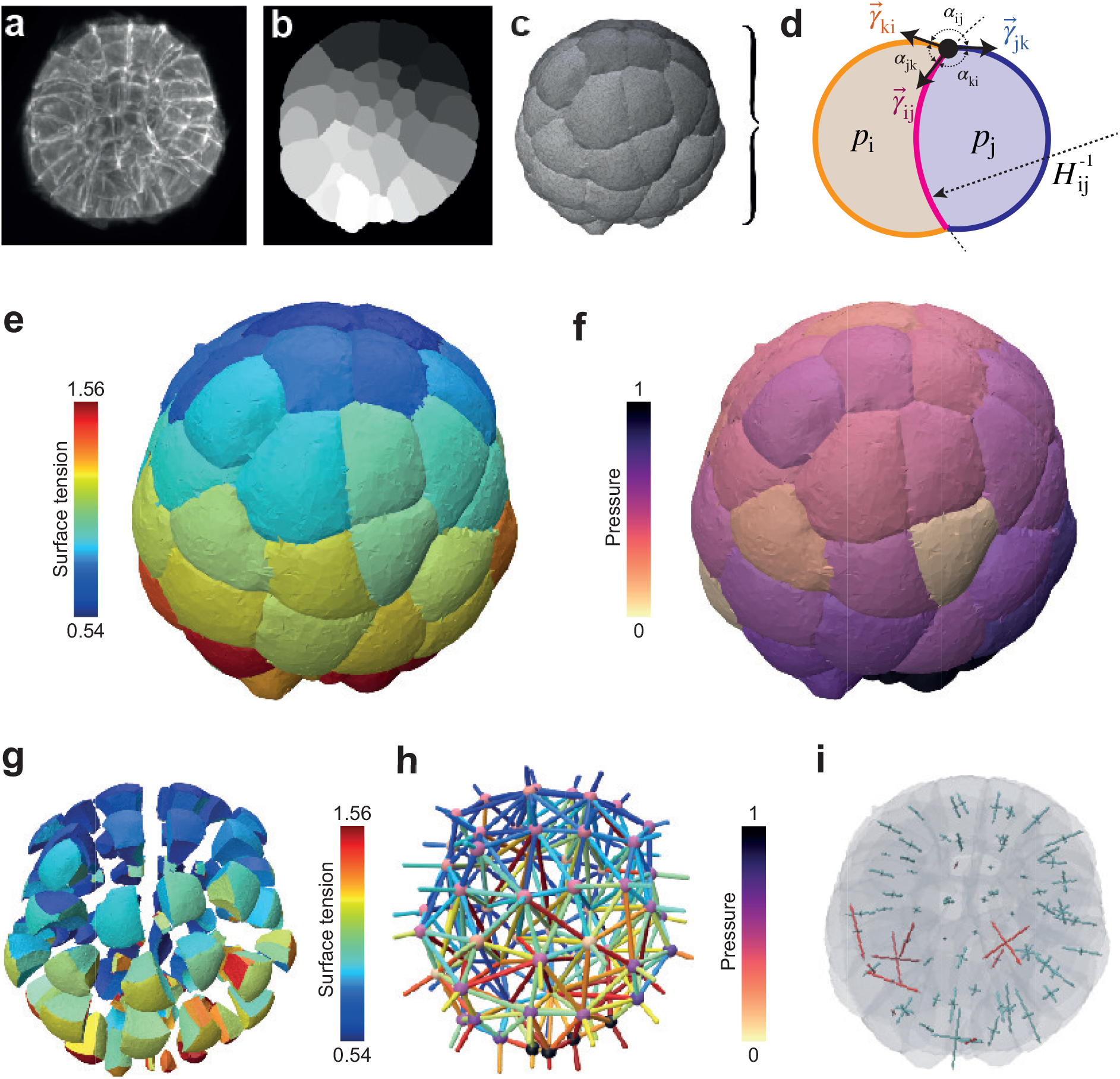
3D force inference procedure and resulting mechanical atlas for a 64 cell ascidian embryo. **a)** 3D fluorescence microscopy image (max projection) of a 64-cell *Phallusia mammillata* embryo (from [23]). **b)** Cell segmentation mask in one focal plane of the 3D image. **c)** Multicellular surface mesh of cell interfaces. **d)** Schematic cell doublet illustrating the two force balances that need to be inverted: the Young-Dupré equation that relates surface tensions γ_*ij*_, γ_*ik*_ and γ_*jk*_ with contact angles *α*_*ij*_, *α*_*ik*_ and *α*_*jk*_, and the Young-Laplace equation that relates cell pressure difference *P*_*j*_ *− P*_*i*_ with tension γ_*ij*_ and the radius of the interface curvature 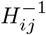. **e)** 3D map of relative surface tensions in the embryo, plotted with a color code from blue (lowest) to red (highest). **f)** Pressure map in the embryo, normalized from 0 to 1. **g)** Exploded view of the surface tension map that illustrates cell-cell contact tensions within the embryo. **h)** Force graph representation of the mechanical atlas, where each node represents a cell with its associated pressure and each edge corresponds to an interface colored by its tension value. **i)** 3D stress eigenvalue representation, corresponding to a stress tensor calculated per cell with the Batchelor formula [49]. Positive eigenvalues are plotted in blue (compressive stress) while negative are plotted in red (extensile stress).

**Fig. 2:**
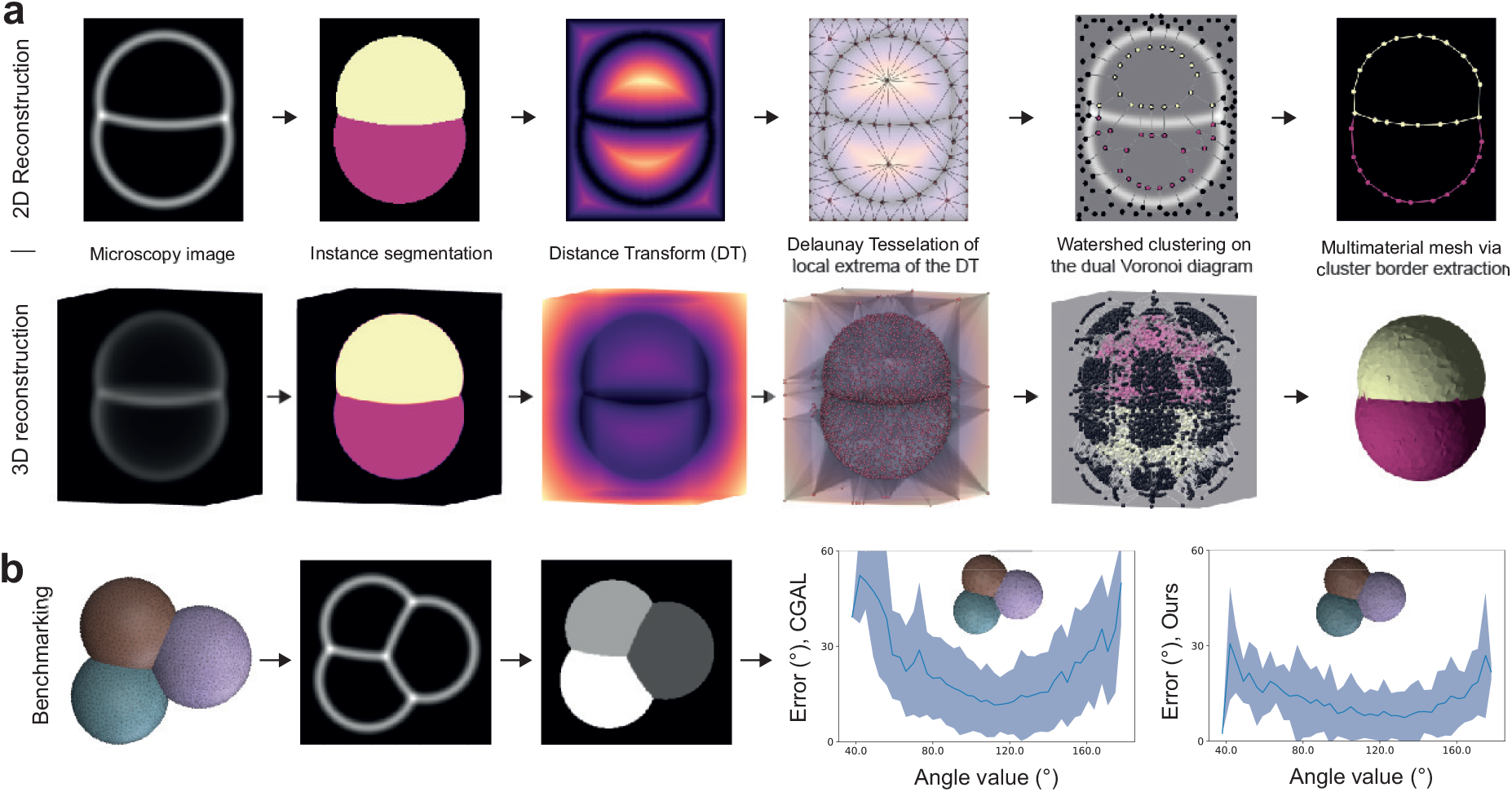
Multimaterial mesh generation algorithm. **a)** From a microscopy image (artificial here) in 2D (resp. 3D), we first generate a distance transform map, including the image boundaries; we then sample points at the extremum values of this map to generate a Delaunay triangulation (resp. tetrahedralization) of the 2D (resp. 3D) space; the average integrated elevation value along edges (resp. triangles) of this tessellation gives weight to edges in the dual Voronoi diagram; a watershed cut algorithm [62] is applied to this weighted graph to partition nodes into cell and exterior regions, resulting *in fine* in a multimaterial nonmanifold polygonal mesh segmentation (resp. triangle surface mesh) of the original cell membrane image. **b)** The geometric precision of our mesh generation algorithm is benchmarked on foam-like simulations, which are transformed into artificial images to reconstruct surface meshes. Our pipeline reconstructs cell geometry with better precision than state-of-the-art mesh generation methods, such as CGAL [57], as shown by the comparison of the error in the reconstructed angles as a function of the original angle. The shaded region measures the standard deviation.

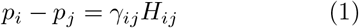

relates the hydrostatic pressure difference *p*_*i*_ − *p*_*j*_ between cells of indices^4^ *i* and *j* with the interface tension 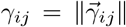 and the interface mean curvature *H*_*ij*_, which is homogeneous along each interface. The Young-Dupré force balance

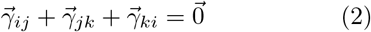

states that the sum of vectorial tensions should be zero at each tri-cellular junction line that joins the interfaces between cells *i, j* and *k*. This vectorial sum is equivalent to saying that tensions are coplanar and form a triangle, which implies the triangle inequality *γ*_*ij*_ < *γ*_*jk*_ + *γ*_*ki*_ and equivalent relations by permutation of the indices *i, j* and *k*. Non-compliance with one of these inequalities indicates that tension balance breaks down and predicts generically a topological transition in the embryo or tissue. The Young-Dupré tension balance can generically be decomposed into a set of two independent scalar equations that combine the polar angles between the interfaces *α*_*ij*_, *α*_*jk*_ and *α*_*ki*_ (Fig. 1d). In the following, we use five different variants of tension balance that involve cosines and sines of polar angles only [30], which we named *Young-Dupré, Young-Dupré projection, Lami, inverse Lami* and *Lami logarithm* [68] (see Methods 5 and Supplementary Note)

The balance of forces in a foam-like tissue of n_*c*_ cells can also be derived from the minimization of surface energy under cell volume constraints. This formulation is particularly adapted to numerical simulations on a discrete mesh [52, 53] and is based on a Lagrangian function. Assigning an index *m* to each existing interface *𝒜*_*m*_ between one cell and another or the external medium, the Lagrangian reads

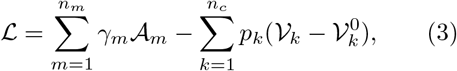

where *γ*_*m*_ and *A*_*m*_ are, respectively, the surface tension and the area of the interface between the regions *a*_*m*_ and *b*_*m*_ where {*a*_*m*_, *b*_*m*_} ∈ ⟦0, *N* ⟧^2^. 𝒱_*k*_ and 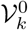 are the current and target volumes in the cell *k*, and *p*_*k*_ its pressure plays the role of a Lagrange multiplier for volume conservation. Discretized on a mesh, where areas and volumes are functions of the positions of the *n*_*v*_ vertices 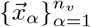 (Extended data Fig.1 and Supplementary Note), optimality conditions [69] for this Lagrangian function produce a force balance at each vertex 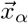

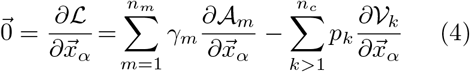

We define Γ = (*γ*_1_, *γ*_2_, …, *γ*_*nm*_)^*T*^ a generalized vector of tensions of size n_*m*_, and 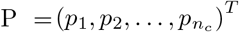 a generalized vector of pressures of size *n*_*c*_. Inspired by projection methods [52] used generically to solve constrained optimization problems, we derive from equation (4) a linear system of equations whose solutions are - to an arbitrary factor - the tensions and pressures corresponding to a given heterogeneous foam geometry (see Supplementary Note). It reads, in matrix form

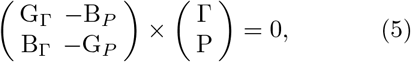

where G_Γ,*P*_ are symmetric matrices of sizes, respectively 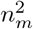 and 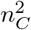 and B_Γ,*P*_ rectangular matrices of sizes, respectively *n*_*m*_ × *n*_*C*_ and *n*_*C*_ × *n*_*m*_ (see Supplementary Note). The linear system to solve for pressures at given tensions G_P_ × *P* = *B*_Γ_Γ is called *Variational Laplace* in the next. Because the square matrix G_*P*_ is full rank and, therefore, invertible (see the Supplementary Note), we can also write a closed-form linear system for the tensions alone as 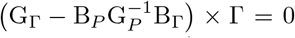, which we call *variational Young-Dupré*.

## 3 Tension and pressure inference

Tensions depend only on contact angles at tri-junctions and are independent of cell pressures, so here we decompose the inverse problem into two steps, in the same spirit as [38]: first, we solve the tensions and then determine the cell pressures using inferred tension values. The advantage of this two-step approach is that tensions can still be inferred in embryos or tissues under confinement or compression (such as *C. elegans*), where Laplace’s force balance does not apply anymore, since the interfaces may adopt non-uniform mean curvatures. Importantly, tensions (and pressures) are known up to a multiplicative (respectively, an additive) factor. To remove this indeterminacy, we impose that the average tensions shall be equal to unity, which adds an equation to the system, and we arbitrarily fix the external pressure to zero. The tension inference problem can be generically cast into a linear system A_Γ_ × Γ = *c*_Γ_, where A_Γ_ is a matrix of size (*n*_Γ_ + 1) × *n*_*m*_ that collects *n*_Γ_ + 1 equations that relate the *n*_*m*_ unknown tensions, and *c*_Γ_ = (0, …, 0, *n*_*m*_)^*T*^ implements the constraint on the average tensions. This system is overdetermined and is solved in the sense of ordinary least-squares (OLS). Performing a systematic benchmark of our method, we found that better results are obtained when the *n*_Γ_ tension equations are weighted by the length of the corresponding junction (see Supplementary Note), which is the choice taken further.

In Fig. 3, we compare the sensitivity of our inference algorithm for the different variants of the Young-Dupré formula (6), (7), (8), and the *variational* Young-Dupré equation. By perturbing vertex positions with random noise in mesh solutions of foam-like simulations (Fig. 3a), we calculate and plot the mean square error on the tensions inferred from this perturbed mesh (Fig. 3b). At low noise values, we find that the scalar Young-Dupré equation gives better results, but this error increases then faster for larger noise. *Variational* Yound-Dupré and the different Lami variants have an error that increases faster at low noise, but then reaches a lower relative plateau at higher noise.

**Fig. 3:**
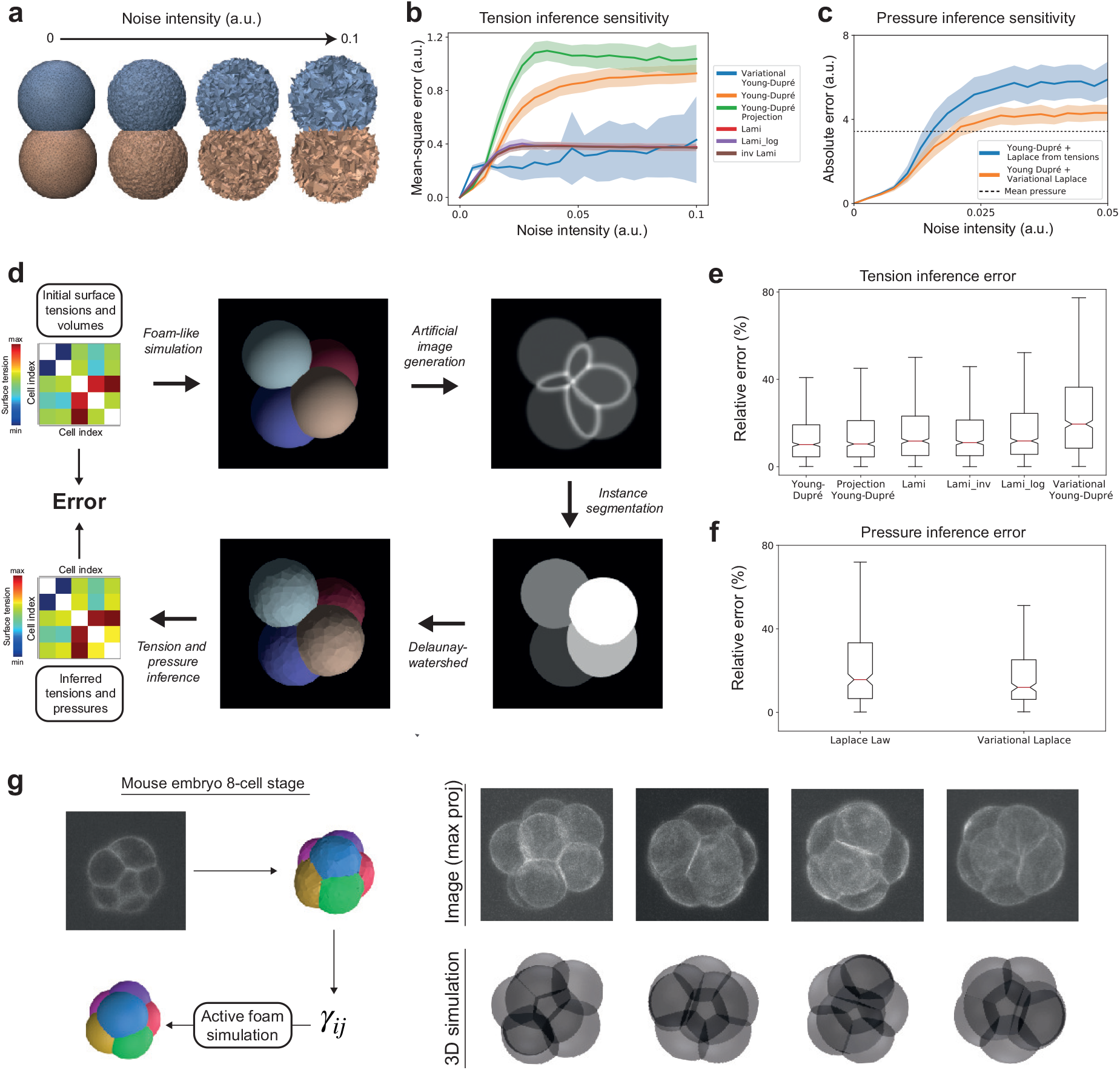
*In silico* validation of the force inference pipeline. **a)** Sensitivity analysis of different formulas for tension and pressure inference. Foam-like simulation meshes are perturbed by randomly displacing each vertex position following a uniform law. **b)** Plot of the mean square error on inferred tensions as a function of the intensity of the noise for the different tension formulas. **c)** Plot of the absolute error on the inferred pressure as a function of the noise intensity for the Laplace and variational Laplace formulas. **d)** Pipeline for benchmarking force inference: from random surface tensions (visualized here as a symmetrical *n*_*c*_ + 1 *× n*_*c*_ + 1 matrix) and cell volume values, a dataset of foam-like embryo meshes is simulated, from which artificial microscopy images are generated; then, our end-to-end pipeline is applied to regenerate a mesh and infer tension and pressure values. **e)** Plot of the relative error in the inferred tensions for the different tension inference formulas applied to our simulated embryo dataset. **f)** Relative errors on inferred pressures on our simulated embryo dataset with Laplace and variational Laplace formulas. **g)** Self-consistent validation of the inference on the compaction of the 8-cell mouse embryo. Surface tensions are inferred with the pipeline and averaged between the cell-medium and cell-cell interfaces. Foam-like simulations are performed using these tensions and yield an *in silico* embryo morphology that is compared to the real embryo image.

For pressure inference, we follow the same approach, expressing the inverse problem as a linear system A_*P*_ × P = *B*_*P*_, which we solve with the OLS method. Here, we compare the traditional Laplace formula (1) and our new *variational* Laplace formula (5). Interestingly, we find that our mesh-based variational formula performs systematically better regardless of the level of noise (Fig. 3c).

Error in inference results may originate from deviations of cells shape from the solution of an heterogeneous foam or from an insufficient image resolution (Extended data Fig. 3c), but they are also the result of an inevitable intrinsic noise generated by our pipeline that comes from the segmentation and meshing operations. To evaluate which formula may be most adapted given this minimal and ineluctable level of noise, we generate ideal artificial confocal microscopy images from mesh results of foam-like simulations (see Supplementary Note). This dataset [66] is used to benchmark our method: the images are segmented using cellpose [58] and translated into multimaterial meshes with our Delaunay-watershed algorithm to ultimately infer tensions and pressures using the various formulas introduced earlier (Fig. 3d). In general, we find that the systematic error induced intrinsically by our pipeline remains very low, with the best inference results obtained with the scalar Young-Dupré formula (6) and the variational Laplace formula (Figs. 3e-f). For all tension and pressure inference examples shown below, we therefore systematically use the scalar Young-Dupré and variational Laplace formula.

## 4 Force inference applied to early embryo development

To validate the biological relevance of our novel force inference pipeline, we inferred 3D mechanical atlases of mouse and ascidian embryos using fluorescent microscopy images of cell membranes. We first study the self-consistency of the heterogeneous foam model in compacting 8-cell mouse embryos. Compaction corresponds to the extension of internal cell contacts that round up the embryo and was shown by micropipette tension measurements [5] to be characterized by a decrease in the ratio 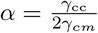 - called *compaction parameter* - where *γ*_cm_ is the tension at the cell medium interface of cells and *γ*_cc_ the tension at cell-cell contacts. This single parameter is enough to characterize the embryo shape and is equal to the cosine of half the contact angle of the cell medium. Using confocal fluorescent images of 8-cell mouse embryos at successive levels of compaction, we segmented them into multimaterial meshes and inferred relative tensions. We then performed 3D foam-like simulations and compared them with the original microscopy images (Fig. 3g), and found a very good qualitative agreement. Interestingly, our automatic inference methods yields systematically lower variability in inferred γ_*cc*_ values than previously obtained by measuring contact angles manually [5], as illustrated on Fig. 3d. This confirms the relevance of a heterogeneous foam model hypothesis and exemplifies the capability of our inference pipeline.

To go beyond this example, where cell-medium and cell-cell tensions are uniform within the embryo, we inferred spatio-temporal mechanical atlases of the early ascidian embryo *Phallusia mammillata*. We used fluorescent images of cell membranes that were acquired with a confocal microscope from the zygote to the 44 cell stage (see Methods) or with a light sheet microscope from the 64 cell stage to the late neurula (≲ 800 cells) [23]. We first focused on the shape of the embryo from 16 cells to the early gastrula, where divisions are reported to be asynchronous with cell divisions that alternate between the animal and vegetal hemispheres [70]. Recently, it was shown in *P. mammillata* embryos at 16, 32 and 44 cell stages, that cells at mitosis entry have lower apical tension than their interphase counterparts located in the opposite hemisphere [6]. This striking result, in notable contrast to mitotic cortical stiffening reported in most somatic cells [71, 72], is again predicted by our force inference method, which finds a ratio of apical tension between mitotic and interphase cells that is systematically lower than 1 in the 16 to 32 cell stages (Fig. 4e). This mitotic softening alternates between the animal and vegetal poles, as illustrated also from pressure maps (Extended data Fig. 4a) further explains the overall 3D shape of the embryo which is flatter on the side of interphase cells (16 and 32 cells). As one would expect from Laplace’s law, if applied globally to the embryo approximated to a droplet, a higher apical tension at one pole leads indeed to its flattening. Inference not only confirms previous results, but also predicts an unknown switch in the 64-cell embryo, where mitotic blastomeres have higher apical tension than their interphase neighbors (Fig 4. 4e, Extended data Fig. 4a) suggesting that, from this stage on, cells undergo mitotic stiffening. This mitotic stiffening persists during gastrulation (stage 120 in Fig. 4e) and later (Extended data Fig. 4a). This illustrates the predictive power of our inference pipeline, which reveals novel mechanical features that explain the shape of cells and embryos.

**Fig. 4:**
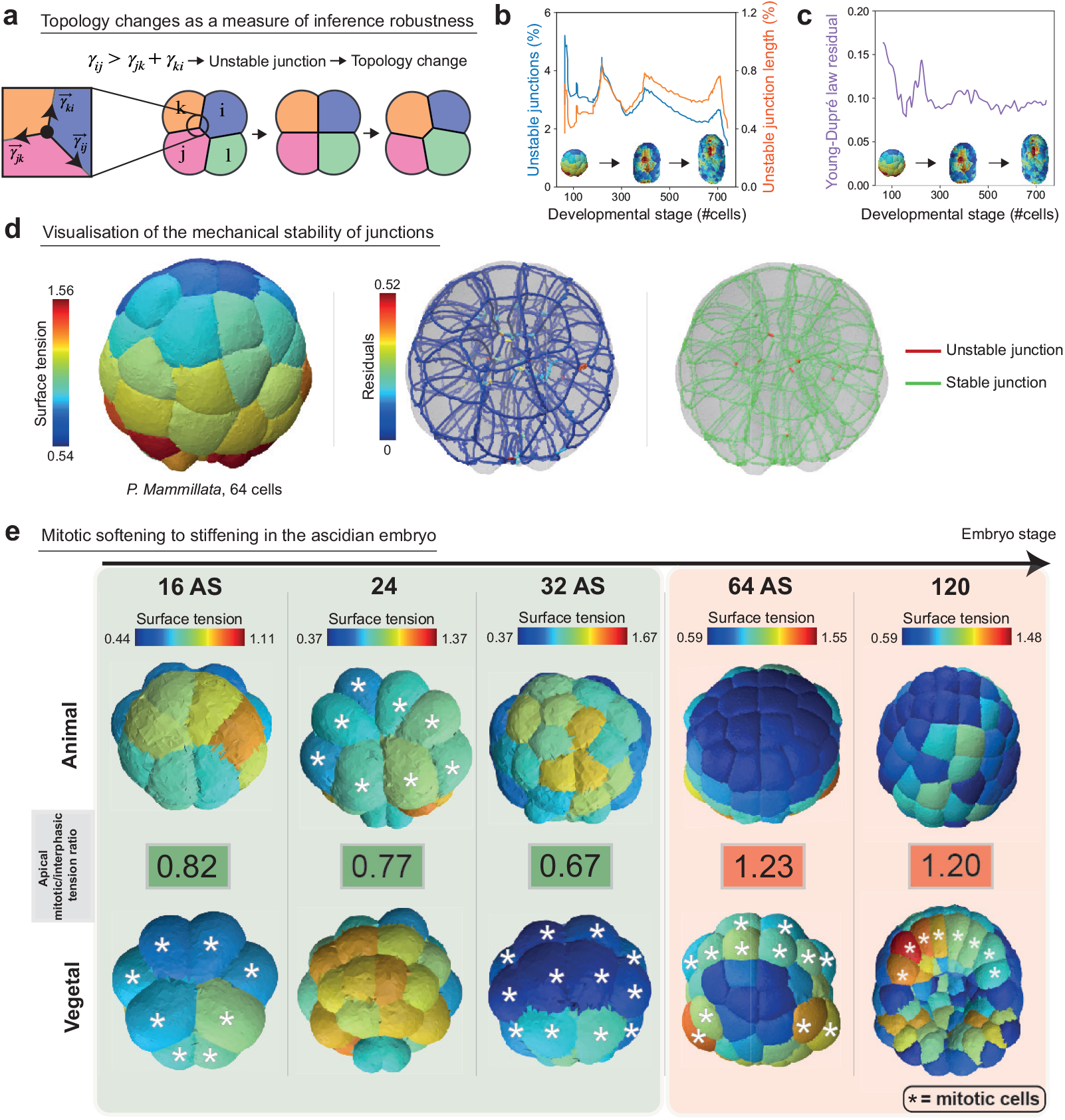
*In vivo* validation of the 3D tension inference. **a)** Illustration of the process of T1 topological transition when one tension at a junction becomes greater than the sum of the two others. **b)**Plot of the percentage of unstable junctions in the embryo (blue) and the ratio of unstable junction length to total junction length in the embryo as a function of its development stage, defined by its number of cells. **c)** Plot of the mean residual of Young-Dupré equations (purple) in the embryo as a function of its development stage, defined by its number of cells. **d)** *Left* Surface tension map of the 64-cell ascidian embryo. *Middle* Visualization of the residuals of Young-Dupré equations for each junction in the same embryo. *Right* Junctions for which inference predicts a T1 topological transition are mechanically unstable. Visualization of stable (green) and unstable (red) junctions in a 64-cell ascidian embryo (*P. mammillata*). **e)** Maps of apical tension at the animal and vegetal poles of the early ascidian embryo (*P. mammillata*) in the 16AS, 24, 32AS, 64 and 120 cell stages. The ratio of mitotic to interphase apical tension is colored green if it is less than 1 and red if it is greater than 1. Mitotic cells are indicated by a white star.

To further assess the validity of our inference method, we searched for locations in the embryo where the hypothesis of foam-like mechanical equilibrium may break down. An interesting idea is to look for junctions that are unstable for the predicted tensions. In fact, when *γ*_*ij*_ > *γ*_*jk*_ + *γ*_*ki*_, we expect the junction ij to be unstable and undergo a T1 topological transition (Fig. 4a). Any unstable junction is therefore the sign of mechanical equilibrium breakdown that can result either 1) from a too large error in tension inference or 2) from an inadequacy of the heterogeneous foam model to describe cell arrangement or geometry [73]. In a 64-cell *P. mammillata* embryo, we found 31 unstable junctions in a total of 569 junctions (Fig. 4d). Interestingly, these unstable junctions are detected exclusively close to the embryo center, where the lengths of the junctions become very small, and segmentation struggles to resolve cell geometry (Fig. 4d, Extended data Fig. 4b). In general, the percentage of unstable junctions predicted by our inference pipeline remains very low, around *≈* 3%, throughout the development of the ascidian embryo up to late neurula (Fig. 4b). This represents an even lower percentage of unstable junction length, below 1%, which confirms that the tension equilibrium predicted by our inference pipeline is generally satisfied. To assess the validity of the inference, it is also useful to visualize the deviation from equilibrium using the force balance at the junctions (2). We therefore propose a visualization of the residuals ∥A_Γ_ × Γ − *c*_Γ_ ∥^2^ at each trijunction, as shown in Fig. 4c,d and Extended data Fig. 4b.

To further illustrate the capabilities of our inference method, we report three aspects of early ascidian embryo development brought to light by our mechanical atlases. The early development of the ascidian is characterized by its high degree of invariance [23], and a stereotypical feature of this invariance is the bilateral symmetry of the embryo. However, each embryo shows a certain degree of geometric variability between its left and right sides, which is well reflected in the mechanical asymmetry, as illustrated by the (a)symmetry of the tension and pressure maps inferred in Fig. 5a and Extended data Fig. 5a.

**Fig. 5:**
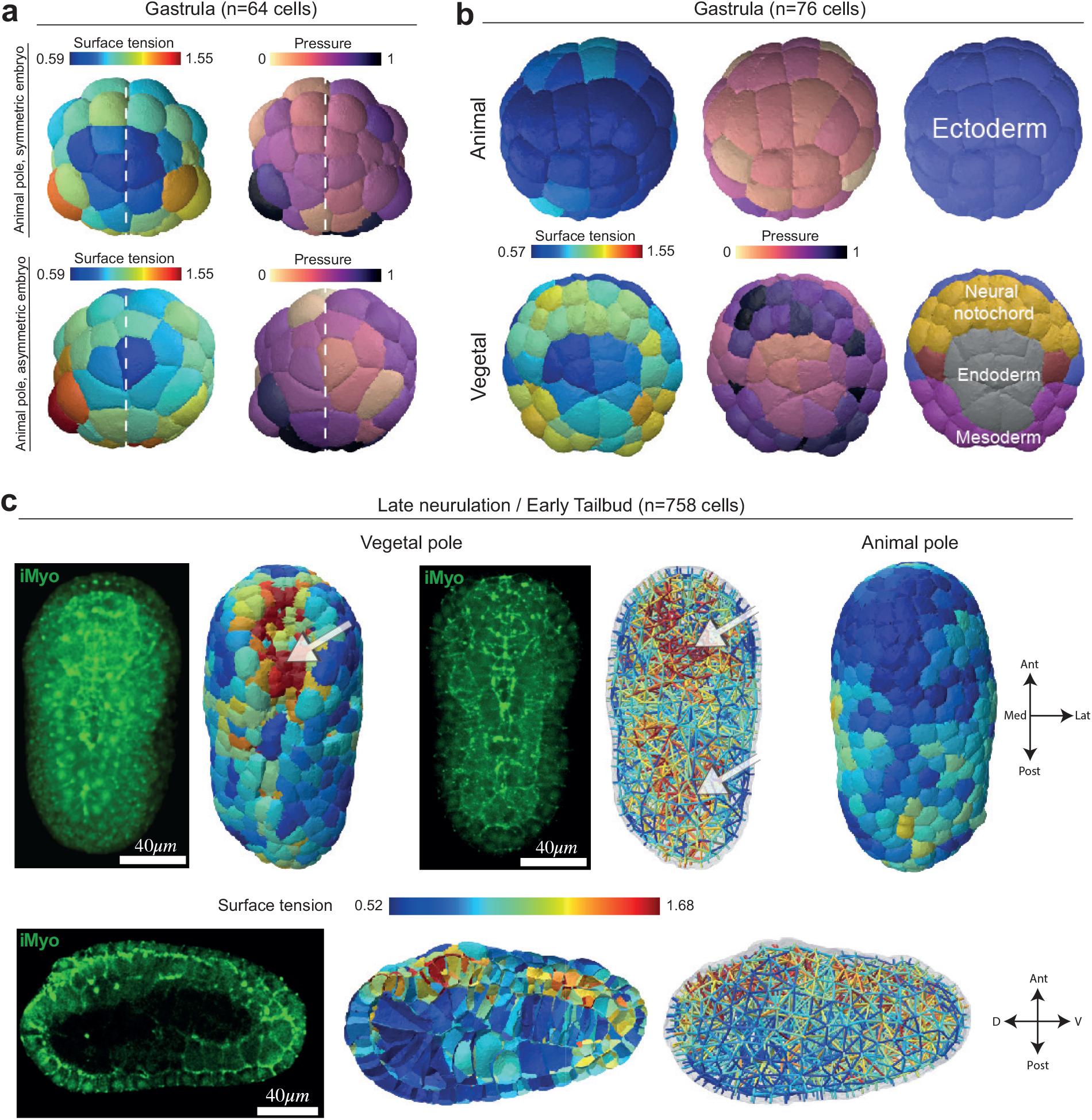
Spatiotemporal patterning of mechanics in the ascidian embryo *P. mammillata*. **a)** Tension and pressure maps of the animal pole of two 64 cell embryos. Imperfections in the geometric bilateral symmetry of the embryo are reflected by a corresponding asymmetry in the apical tension and pressure of the cell. **b)** Tension and pressure maps at the animal and vegetal poles of a 76-cell embryo and the corresponding pattern of cell fate in the germ layers. **c)** Tension maps in late neurula (758 cells) from vegetal, animal, and sagittal views. The white arrows indicate regions of higher tension (red) within the embryo. Fluorescent microscopy images of myosin II (iMyo) at the vegetal pole and in the sagittal view: 3D reconstruction (top left) or selective plane projection of 10 confocal planes (top middle and bottom left). The orientation of the embryo is given by arrows Ant: anterior, Pos: posterior, Med: medial, Lat: lateral, D: dorsal, V: ventral.

The cell fate in the ascidian embryo is also invariant, as has been described for several decades (reviewed in [23, 74]). At the 76-cell stage, the animal hemisphere is composed exclusively of ectodermal cells, while the vegetal hemisphere is segregated into neural/notochord progenitors and endoderm or mesoderm germ layers. We find that this patterning of cell fate is reflected in a remarkable manner in different regions of cell mechanics: ectoderm and endoderm cells have lower apical tension and lower pressure, while neural plate and mesoderm cells form very distinct regions of higher apical tension and pressure (Fig. 5b). This is probably due to the different mitotic history of each lineage, since fate specification is accompanied by an independent cell cycle timing in each specified tissue [23, 70]. In the 76-cell stage, neural/notochord and mesoderm cells have, in fact, just undergone cell division (they are in their eighth cell cycle), while endoderm cells were born more than 40 minutes ago and are in the middle of interphase, just before they undergo apical constriction [75]. In the neurula stage, apical constriction has been reported to drive neural tube closure with greater contractility on the apical side of the nerve cord and brain tissues [76, 77]. Consistent with this, our inference pipeline predicts on the vegetal side of the embryo at 395, 702 and 758-cell stage a high apical tension in cells located in the anterior neural plate that are undergoing folding (Fig. 5c arrow in the vegetal pole view, Extended data Fig. 5b). A sagittal section of the embryo at this stage reveals that the neural tube has more cortical tension than the overlying epidermis of the underlying endoderm and notochord (Fig. 5c sagittal section); this higher tension is reflected in a stronger accumulation of myosin II in the neural tube compared to other tissues (Fig. 5c, myosin sagittal section and see also [76]).

Finally, we performed tension inference in the early *C. elegans* embryo from 4 to 15 cells (Fig. 6). Unlike ascidian and, to a certain extent, mouse embryos, an eggshell strongly constrains the shape of cells from the zygote stage. This confinement has shown to be an essential cue controlling early cell arrangement [78, 79] and makes Laplace’s law no longer adequate to account for cell pressures, which are directly affected by the mechanical resistance of the shell. We confirm this characteristic with 3D simulations of a 4 cell embryo confined within an ellipsoid (Fig. 6a), using realistic parameters that we previously measured in [79]. In this realistic simulation, we show that the mean curvature may be locally perturbed by the shell along cell-medium interfaces, especially for ABp and EMS blastomeres, which precludes the use of Laplace’s law, which assumes constant mean curvature interfaces. Indeed, when we infer pressures with the Laplace or Laplace variational formula on this mesh, we obtain pressure predictions, which are 20% to 30% different from the actual value in the four blastomeres (Fig. 6a). Therefore, simultaneous tension and pressure inference may not be a good strategy in this case [41], while breaking down the inference in two successive steps still allows us to infer tensions independently of cell pressures. Interestingly, we find, in agreement with the measurements in [79], a lower cell-medium tensions in P2 and EMS cells in the 4-cell stage *C. elegans* embryo, and predict a general trend of lower cell-medium cortical tension in descendants of the P-lineage at subsequent stages of embryo development (Fig. 6b).

**Fig. 6:**
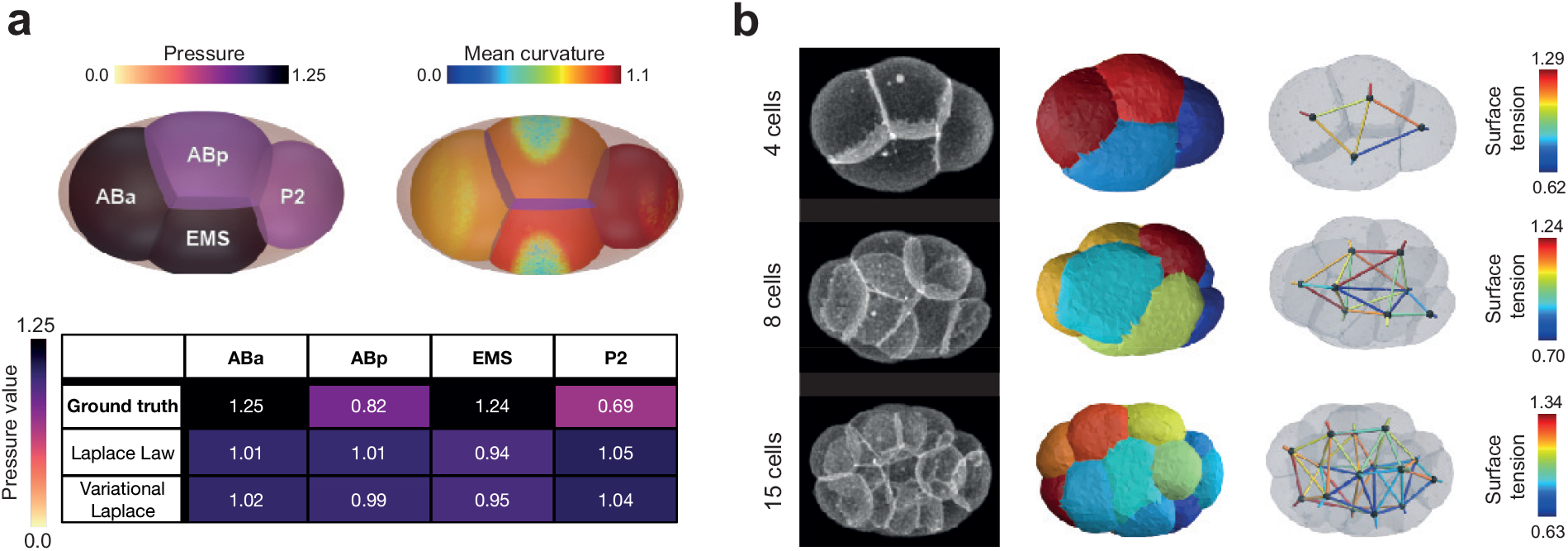
Force inference in *C*.*elegans* embryo. **a)** Pressure inference on a simulated embryo confined in a rigid shell. The shell induces deformations in the membrane that lead to spatial changes in curvatures compared to those of an isolated foam. Both the Laplace and the variational Laplace formulas are inadequate to infer correct pressures, as illustrated by the table of values. **b)** Surface tension can still be inferred, as equilibrium at junctions is still verified.

## 5 Discussion

We presented a robust end-to-end computational pipeline to infer relative surface tensions and pressures directly from three-dimensional fluorescent images of embryos or tissues. It is based, in particular, on a novel and fast method for generating surface meshes from cell segmentation masks, which allows for a more accurate extraction of geometric features than previous approaches [57]. Therefore, our algorithm is compatible with the latest segmentation methods [58, 80–82] and can scale to thousands of cells. We also introduced a novel formula for inferring pressures from a triangle surface mesh, which outperforms the direct inversion of Laplace’s law. By performing a systematic sensitivity analysis on simulated embryos, we showed that the classic Young-Dupré formula gives the best tension inference results for moderate noise in the image or in the cell shape. Our pipeline intrinsically achieves maximum relative force errors of *≈* 10% from images of simulated embryos (Fig. 3c and Supplementary Note). Additionally, we provide several visualization tools to display multicellular morphology and forces in multiple ways, including a force graph representation of the cell aggregate and a 3D map of cellular stress tensors (Figs. 1h-i). The residues and predicted topological changes of inference for each junction in the aggregate can also be directly plotted to enable local evaluation of the method and/or the active foam hypothesis (Fig. 4b,c). Subsequently, we demonstrated the biological relevance of our approach by generating mechanical atlases of the early ascidian embryo: our inference method can recover characteristic patterns of apical tension previously observed [76], including a lower apical tension measured in mitotic cells before 64-cell stage [6]. Interestingly, it can also make new predictions and reveal mirroring patterns of cell mechanics and cell fate in germ layers. Finally, we demonstrate the utility of decoupling pressure and tension inference by applying our methodology to the early *C*.*elegans* embryo, which develops within a shell.

One forthcoming challenge will be to generate spatio-temporal mechanical atlases of various embryos. Indeed, a temporal reference is so far missing to calibrate the successive spatial maps in time. As demonstrated in 2D [79], combining static inference with the temporal measurement of absolute forces in a single location, or imaging phosphomyosin fluorescence intensity as a proxy for tension, could become a generic approach to construct temporal atlases of absolute mechanical forces, but this needs to be repeated in 3D.

A second challenge will involve the inclusion of junctional mechanics in the form of additional line tension contributions at the apical surface of cells. Indeed, blastomeres with a contact to the cell medium acquire generally apico-basal polarity short before the blastula stage in early embryos. This emergence of apical polarity is generally associated with the formation of tight junctions and a contractile ring of actomyosin delimiting each apical surface [83, 84], that is expected to create additional line tensions at tricellular junctions. The question of the uniqueness of the inverse solution will furthermore arise, since several stable discontinuous bifurcation states can exist in the presence of line and surface tensions [85, 86], which will first require a in-depth theoretical effort.

A third challenge will consist of generalizing force inference methods to more complex mechanical models, such as recent active viscous surface models [54, 85, 87–90], which naturally generate inhomogeneous and anisotropic surface tensions, as well as possible torques, leading to more complex shapes and force balance equations. This will be particularly important for precisely characterizing the mechanics of dividing cells and faster growing organisms, such as *C. elegans*, for which the time scales of visco-active relaxation and development may no longer be well separated. A possible generic avenue to solve these problems may lie in a fully variational approach, where a mathematical loss between the microscopy images and the meshes could be constrained by an arbitrary mechanical model to allow direct gradient-based optimization of its spatio-temporal parameters. Our recent effort to design such an efficient loss for comparing a mesh and an image may begin to fill this gap [91]. Importantly, the current force inference method we introduced will remain a fundamental building block to this research field, providing already accurate geometric and mechanical maps, which will form an ideal initial guess to refined but more computationally expensive iterative methods.

With a documented and user-friendly implementation in Python [48], our 3D force inference method can be easily applied to 3D images of embryos or small tissues undergoing a sufficiently slow development, and can be combined with spatial ”omic” data generated in early embryos to uncover possible mechanochemical couplings. 3D force inference complements the growing range of tools available for studying the mechanical properties of tissues in space and time [25, 92, 93], and we anticipate that this approach will help elucidate the mechanical underpinnings of large-scale morphogenetic movements at the cellular level and illuminate the intricate interplay between chemical signaling and mechanics during development [94–96]. By revealing the developmental forces shaping organisms, our method may open new evo-devo studies, such as the investigation of the mechanical differences between closely related phylogenetic neighbors or the understanding of the mechanical aspects contributing to the divergence of developmental pathways in evolution.

## Data availability

Images and segmentation masks are already available publicly for *P. Mammillata* embryos on https://figshare.com/projects/Phallusia_mammillata_embryonic_development/64301 (*≥* 64 cells) [23] and for *C. elegans* embryos on https://doi.org/10.6084/m9.figshare.12839315[22].

The simulated dataset (original simulation meshes, artificial images, segmentation masks and tensions/pressures) used to benchmark the method is available publicly on https://doi.org/10.5281/zenodo.7881017[66].

Additional experimental images of ascidian embryos (< 64 cells) and their segmentation masks are available upon request.

## Code availability

Our inference pipeline *foambryo* [48] is distributed as a standalone Python package in the repository PyPI and the source code is available on GitHub and is archived on Zenodo. The mesh reconstruction pipeline *Delaunay-watershed* [67] is also distributed as a separate Python package on PyPI and its source code is on GitHub and archived on Zenodo.

## Acknowledgments

This project has received funding from the European Research Council (ERC) under the Horizon 2020 research and innovation program of the European Union (Grant agreement No. 949267). SI was funded by Ecole Polytechnique (AMX grant). HT has been supported by EMBRC-France (AAP Découverte 2020), by the Bettencourt-Schueller Foundation, by the CNRS and the Collège de France. RD and AMD are supported by a grant from the French Government funding agency Agence Nationale de la Recherche to McDougall (ANR ‘MorCell’: ANR-17-CE13-0028) and received funding from the MITI at CNRS (AAP Modélisation du vivant). The authors are grateful for continuous support to the Imaging Platform (PIM) and the Animal Facility (CRB) of the Institut de la Mer de Villefranche (IMEV), which is supported by EMBRC-France, whose French state funds are managed by the ANR within the Investments of the Future program under reference ANR-10-INBS-0. The authors thank J-L. Maître for sharing microscopy images of 8-cell-stage mouse embryos, and F. Graner as well as all members of the Turlier team for discussions.

## Authors’ contributions

H.T. supervised the project and acquired funding. S.I., F.D. and H.T. developed the theory. S.I. designed the computational pipeline and performed all computations. R.D. and A.D. performed the experiments. S.I., R.D. and H.D. analyzed the data. S.I. made the figures with input from H.T.. H.T. wrote the manuscript with the input of all authors.

## Competing interests

The authors declare that they have no competing interests.

## Open Access

For the purpose of Open Access, the author has applied a CC BY 4.0 public copyright license to any Author Accepted Manuscript version arising from this submission. The codes and softwares of this manuscript are available in open-source and licensed under CC BY-NC 4.0.

## Methods

### Variants of Young-Dupré formulas

Starting from the vectorial expression of the Young-Dupré law (2) we call its decomposition simply by *Young-Dupré* its decomposition with cosines of polar angles:

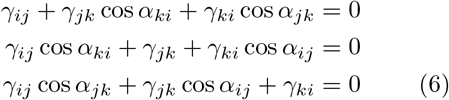

Another set involves both cosines and sines of angles made by vectorial tensions with one direction chosen arbitrarily choose along a tension vector, and we call it *Young-Dupré projection*:

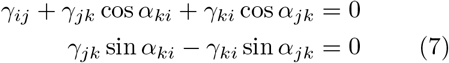

Many other mathematically equivalent formulas may in fact be derived from trigonometric laws applied to the triangle formed by vectorial tensions (see Supplementary Note). Here, we will also use Lami’s theorem, which derives directly from the law of sines and was proposed as an alternative formula for tension inference in 2D [42, 68]:

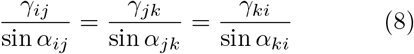

To avoid divergence at small polar angles, it was proposed to consider the same equations written as *γ*_*ij*_ sin *α*_*jk*_ = *γ*_*jk*_ sin *α*_*ij*_, *γ*_*jk*_ sin *α*_*ki*_ = *γ*_*ki*_ sin *α*_*jk*_, which we call *inverse Lami*, or to consider the logarithm of the equation (8), that we call *Lami logarithm*.

### Biological material

The eggs of the ascidian *Phallusia mammillata* were harvested from animals obtained in Sète and kept in the laboratory in a tank of natural seawater at 16°C. Egg preparation and microinjection have been previously described (see detailed protocols in [97], [98]). Eggs and sperm were collected by dissection. Sperm was activated in pH 9.0 seawater prior to fertilization (see the detailed protocol in [98]). All imaging experiments were performed at 20°C.

### Plasma membrane and myosin-II fluorescent labeling

The plasma membrane was imaged using our characterized construct PH::Tomato [98] whereas Myosin II was imaged using Myosin II intrabody iMyo (called SF9::GFP in Chaigne et al., 2016, the plasmid pRN3-SF9-GFP is a kind gift from the M.H. Verlhac laboratory). RNAs coding for PH::Tomato (1 µg.µL-1) and SF9/iMyo::GFP (4 µg.µL-1) were injected in unfertilized Phallusia oocytes that were then fertilized between 2 and 12 hours after injection.

### Confocal imaging of *Phallusia mammillata* embryos

4D confocal imaging was performed at 20 ° C using a Leica TCS SP8 inverted microscope equipped with hybrid detectors and a 20×/0.8NA water objective lens. A 3D stack was taken every minute with a pixel size of 1µm x 1µm and a z-step of 1 µm (in order to obtain cubic voxels). The Phallusia embryos shown in Fig. 4d (and Extended data Fig. 4) from 16 cells to 32 cells were imaged in the Team ABC laboratory, while embryos from stage 64 cells and later stages (shown in Fig. 1, Fig. 4, Fig. 5, Ext Fig. 4, Ext Fig. 5) were obtained from a public dataset of segmented *P. mammillata* embryos published in [23].

### Statistical Analysis

The boxplots (shown in Fig. 3e, Fig. 3f, Ext Fig. 3b) are realized with the default parameters of the boxplot function of the matplotlib python library. The box center is located at the median, and its extremities represents the first and third quartiles. The whiskers are located at Q1 - 1.5*(Q3-Q1) and Q3 + 1.5*(Q3-Q1).

The shaded regions in plots displays the standard deviation (in Fig. 3b, Fig. 3c, Fig. 2b, Ext Fig. 2b).

**Extended data Fig. 1:**
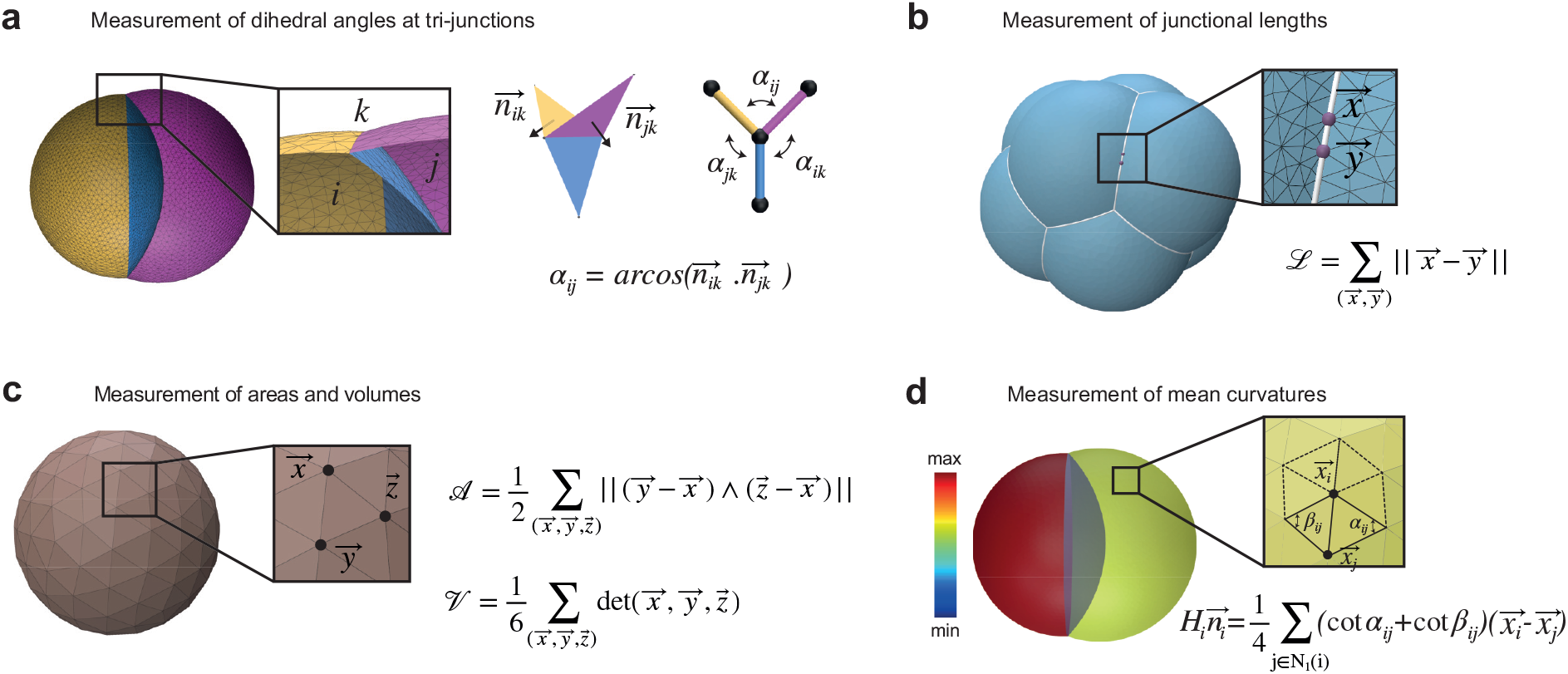
Measurement of geometrical quantities on nonmanifold multimaterial triangle surface meshes. **a)** Contact angles are calculated at each junction as the mean of dihedral angles in each triplet of triangles that constitutes the junction. A dihedral angle is computed from the unit normals to the two adjacent triangles. **b)** Junctions are lines that separate three different materials or regions (three cells or 2 cells and the cell medium). Their length can be easily defined and measured with our nonmanifold mesh data structure. **c)** Each cell is represented by a bounded volume (a discrete manifold). We can compute their volumes and areas from our multimaterial mesh data structure with formulas derived in the Supplementary Note. **d)** Mean discrete curvatures can be computed using the cotangent formula (see Supplementary Note).

**Extended data Fig. 2:**
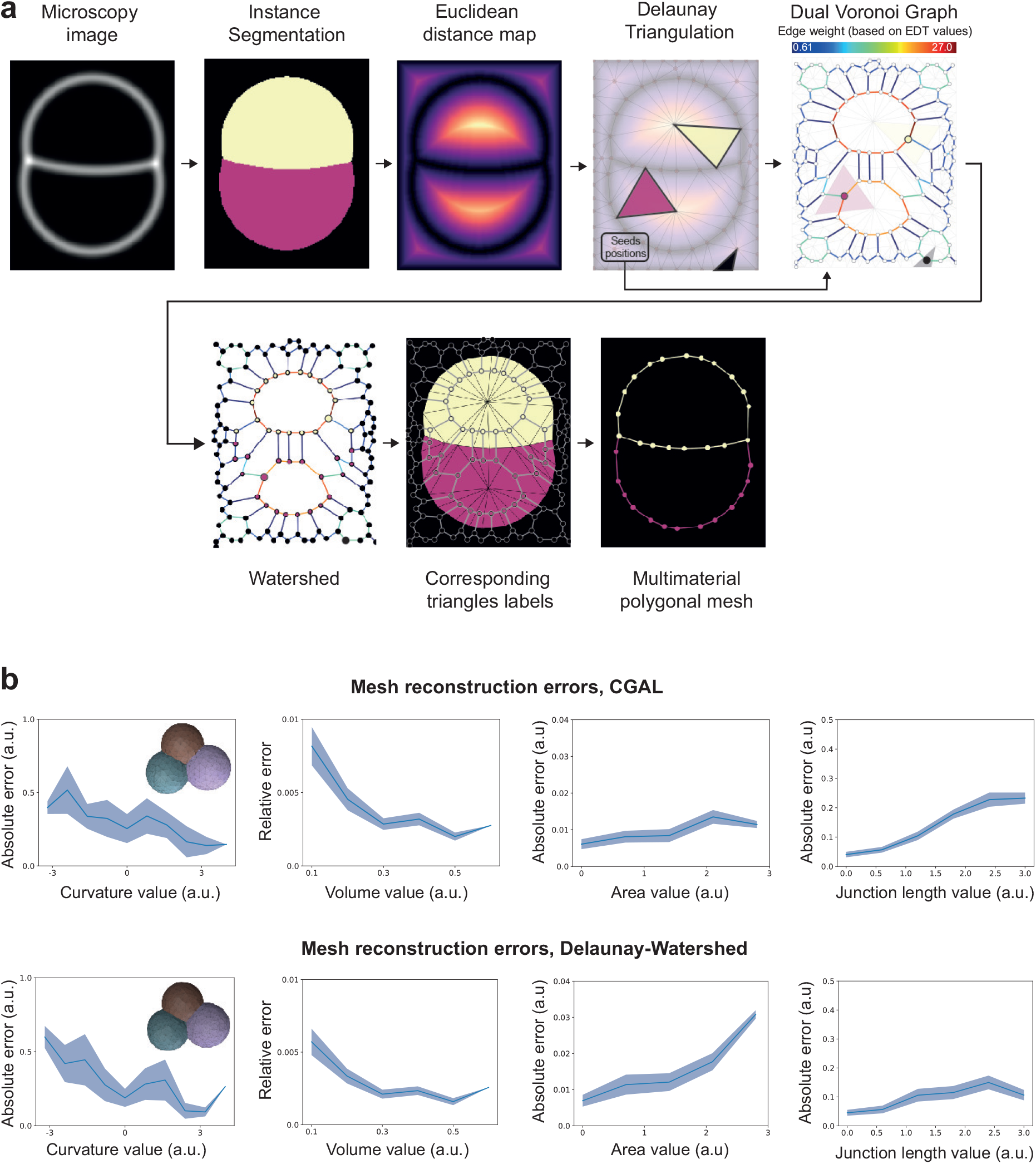
Detailed procedure and benchmarking of the Delaunay-watershed mesh generation algorithm. **a)** Pipeline for mesh generation from a microscopy image (here in 2D for graphical purposes). From the Delaunay triangulation of the image domain, we construct a graph of the dual Voronoi diagram. The edge weights of this graph are computed by integrating the value of the Euclidean distance map along corresponding edges that separates two triangles in the primary domain. The watershed is performed on the dual graph, and the seeds are chosen by taking the triangles containing the pixel with the highest EDT value in the primary domain. **b)** Comparison of the geometric error obtained on interface curvatures, cell volumes, interface areas, and junctional lengths with CGAL and our Delaunay-watershed algorithms for mesh reconstruction.

**Extended data Fig. 3:**
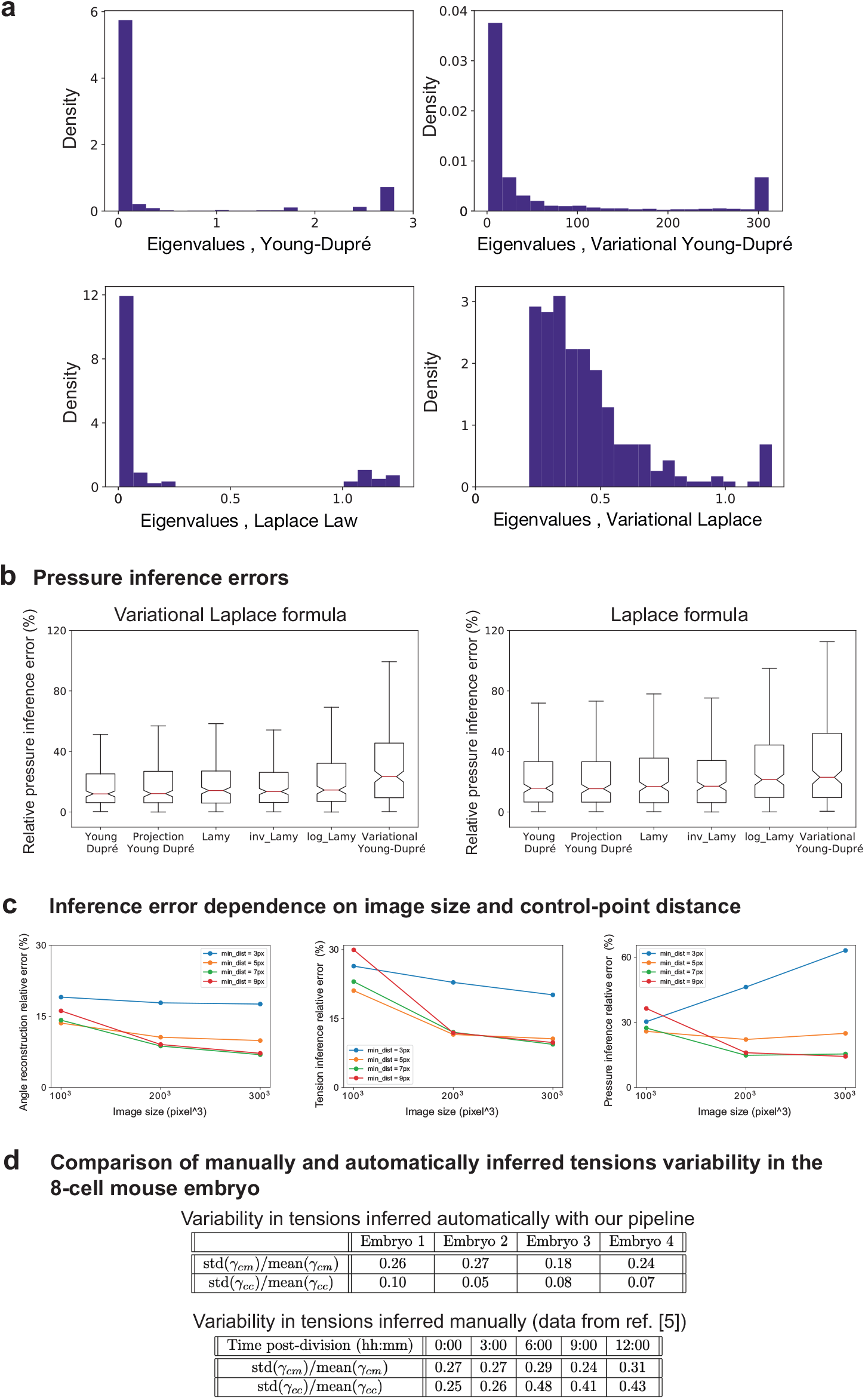
Inference sensitivity and influence of tensions formulas for pressure inference. **a)** Histogram of eigenvalues of the pseudo-inverse matrices used to infer tensions and pressures for the Young-Dupré, variational Young-Dupré, Laplace and Variational Laplace formulas, on our simulated embryo dataset. The spread of the histogram is a measure of the conditioning of the matrix. **b)** Comparison of the relative error on inferred pressures obtained on our simulated embryo dataset between Laplace and variational Laplace formulas. **c)** Mean relative error on angles reconstruction (left), tension inference (middle) and pressure inference (right), depending on the refinement of the mesh (in pixels) and the image size.

**Extended data Fig. 4:**
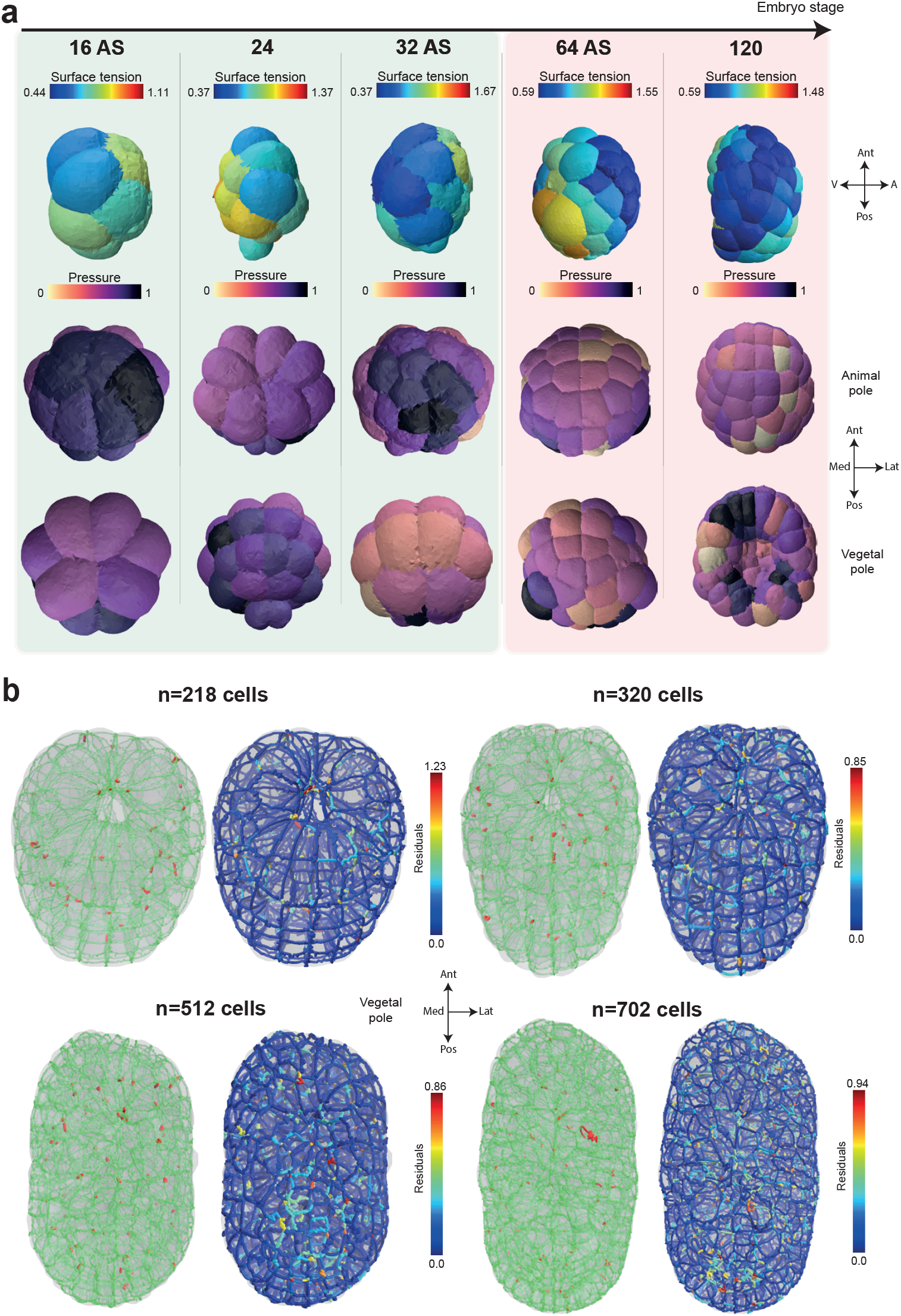
Additional validation data of the 3D tension inference. **a)** Mitotic softening and stiffening in the 16AS, 24, 32AS, 64 and 120 cell stages of the early ascidian embryo (*P. mammillata*). Upper row: sagittal view of inferred apical tension. Middle and bottow rows: animal and vegetal views of the inferred cell pressures. The ratio of mitotic to interphase apical tension is colored green if it is less than 1 and red if it is greater than 1. The orientation of the embryo is given by arrows Ant: anterior, Pos: posterior, Med: medial, Lat: lateral, V: vegetal, A: animal. **b)** Vegetal view of stable (green) and unstable (red) junctions (Left) and tension inference residues (Right) in ascidian embryos (*P. mammillata*) at 218, 320, 512 and 702 cell stages. The orientation of the embryo is given by arrows Ant: anterior, Pos: posterior, Med: medial, Lat: lateral.

**Extended data Fig. 5:**
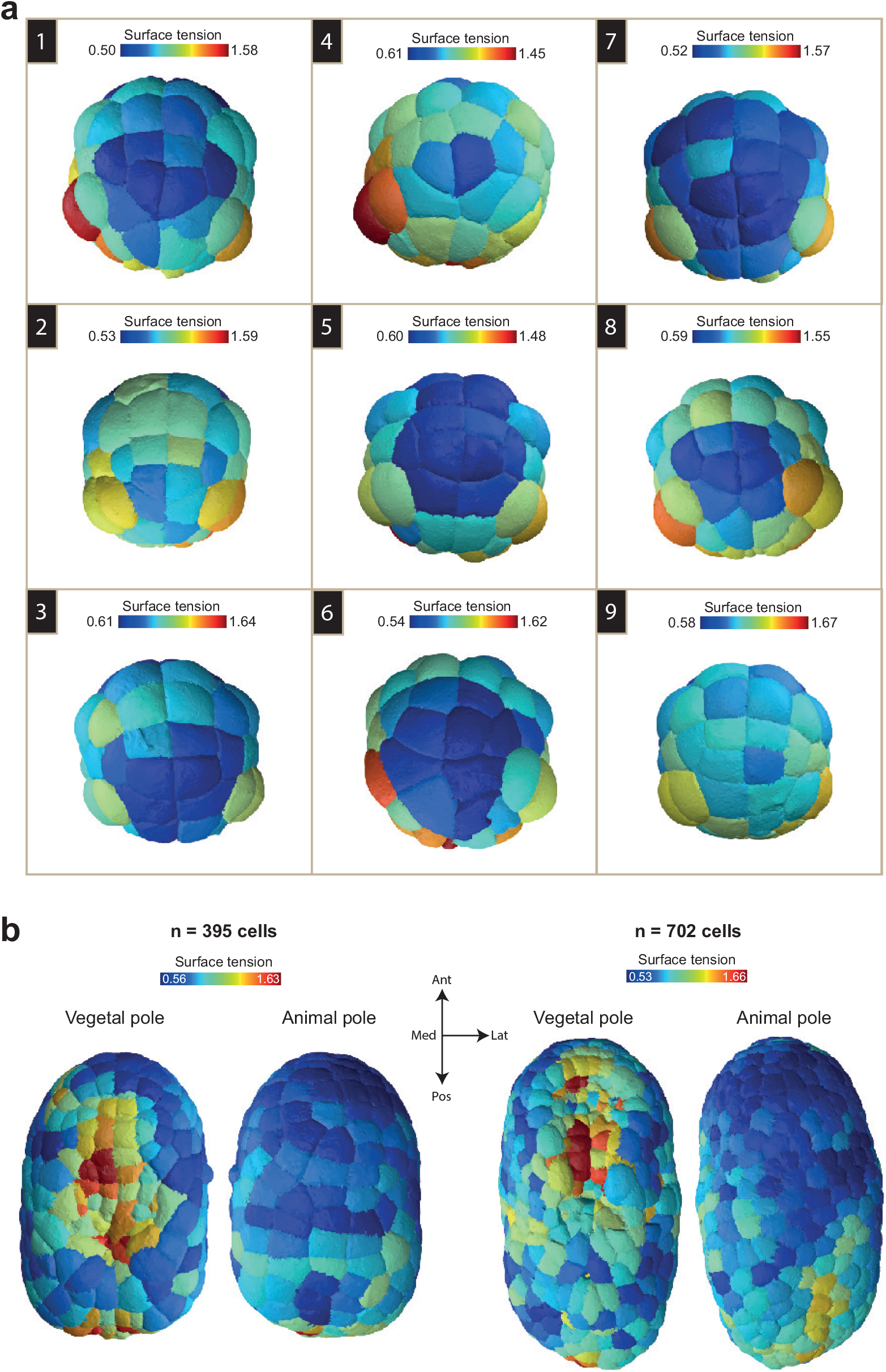
Additional tension maps of ascidian *P. mammillata* gastrula and neurula. **a)** Nine examples of apical tension maps of 64 cell gastrula (animal pole). **b)** Maps of apical tensions at the animal and vegetal poles of early (Left) and late neurula ascidian embryos. The orientation of the embryo is given by arrows Ant: anterior, Pos: posterior, Med: medial, Lat: lateral.

## Footnotes

1 in this paper we used either the deep-learning tool *cellpose* [58] or preexisting segmentation masks

2 This EDT map may also be predicted directly from raw fluorescent images by training a convolutional-neural network [60, 61]

3 Other graph partitioning methods such as multicut [63], hierarchical agglomeration [64] or Mutex watershed [65] algorithms may also be envisioned, although we have not tried them directly.

4 by convention the index 0 will refer to the external medium

## Notes

https://github.com/VirtualEmbryo/foambryo

### Competing Interest Statement

The authors have declared no competing interest.

### Summary of Updates

Figure 4 updated; Extended Figure 3 updated; Main text revised; Supplementary note revised; Repository link to source code added; Typos corrected

## References

[1] Gawad, C., Koh, W. & Quake, S. R. Single-cell genome sequencing: current state of the science. Nature Reviews Genetics 17 (3), 175–188 (2016).

[2] Hwang, B., Lee, J. H. & Bang, D. Single-cell rna sequencing technologies and bioinformatics pipelines. Experimental & molecular medicine 50 (8), 1–14 (2018).

[3] Mitchison, J. & Swann, M. The mechanical properties of the cell surface. J. exp. Biol 31 (3), 443–460 (1954).

[4] Guevorkian, K., Colbert, M.-J., Durth, M., Dufour, S. & Brochard-Wyart, F. Aspiration of biological viscoelastic drops. Physical review letters 104 (21), 218101 (2010).

[5] Maître, J.-L., Niwayama, R., Turlier, H., Nédélec, F. & Hiiragi, T. Pulsatile cell-autonomous contractility drives compaction in the mouse embryo. Nature cell biology 17 (7), 849–855 (2015).

[6] Godard, B. G. et al. Apical relaxation during mitotic rounding promotes tension-oriented cell division. Developmental Cell 55 (6), 695–706 (2020).

[7] Svoboda, K. & Block, S. M. Biological applications of optical forces. Annual review of biophysics and biomolecular structure 23 (1), 247–285 (1994).

[8] Tanase, M., Biais, N. & Sheetz, M. Magnetic tweezers in cell biology. Methods in cell biology 83, 473–493 (2007).

[9] Bambardekar, K., Clément, R., Blanc, O., Chardès, C. & Lenne, P.-F. Direct laser manipulation reveals the mechanics of cell contacts in vivo. Proceedings of the National Academy of Sciences 112 (5), 1416–1421 (2015).

[10] Rheinlaender, J. et al. Cortical cell stiffness is independent of substrate mechanics. Nature materials 19 (9), 1019–1025 (2020).

[11] Fujii, Y. et al. Spatiotemporal dynamics of single cell stiffness in the early developing ascidian chordate embryo. Communications biology 4 (1), 1–12 (2021).

[12] Campàs, O. et al. Quantifying cell-generated mechanical forces within living embryonic tissues. Nature methods 11 (2), 183–189 (2014)

[13] Serwane, F. et al. In vivo quantification of spatially varying mechanical properties in developing tissues. Nature methods 14 (2), 181–186 (2017).

[14] Souchaud, A. et al. Live 3d imaging and mapping of shear stresses within tissues using incompressible elastic beads. Development 149 (4), dev199765 (2022).

[15] Marín-Llauradó, A. et al. Mapping mechanical stress in curved epithelia of designed size and shape. bioRxiv 2022–05 (2022).

[16] Beloussov, L. V., Dorfman, J. G. & Cherdantzev, V. G. Mechanical stresses and morphological patterns in amphibian embryos. Journal of embryology and experimental morphology 34 (3), 559–574 (1975).

[17] Rauzi, M., Verant, P., Lecuit, T. & Lenne, P.-F. Nature and anisotropy of cortical forces orienting drosophila tissue morphogenesis. Nature cell biology 10 (12), 1401–1410 (2008).

[18] Norotte, C., Marga, F., Neagu, A., Kosztin & Forgacs, G. Experimental evaluation of apparent tissue surface tension based on the exact solution of the laplace equation. EPL (Europhysics Letters) 81 (4), 46003 (2008).

[19] Forgacs, G., Foty, R. A., Shafrir, Y. & Steinberg, M. S. Viscoelastic properties of living embryonic tissues: a quantitative study. Biophysical journal 74 (5), 2227–2234 (1998)

[20] Mazuel, F. et al. Magnetic flattening of stem-cell spheroids indicates a size-dependent elastocapillary transition. Physical review letters 114 (9), 098105 (2015).

[21] Mary, G. et al. All-in-one rheometry and nonlinear rheology of multicellular aggregates. Physical Review E 105 (5), 054407 (2022).

[22] Cao, J. et al. Establishment of a morphological atlas of the caenorhabditis elegans embryo using deep-learning-based 4d segmentation. Nature communications 11 (1), 1–14 (2020).

[23] Guignard, L. et al. Contact area–dependent cell communication and the morphological invariance of ascidian embryogenesis. Science 369 (6500), eaar5663 (2020).

[24] McDole, K. et al. In toto imaging and reconstruction of post-implantation mouse development at the single-cell level. Cell 175 (3), 859–876 (2018).

[25] Prevedel, R., Diz-Muñoz, A., Ruocco, G. & Antonacci, G. Brillouin microscopy: an emerging tool for mechanobiology. Nature methods 16 (10), 969–977 (2019).

[26] Bevilacqua, C. et al. High-resolution linescan Brillouin microscopy for live imaging of mechanical properties during embryo development. Nature Methods (2023).

[27] Selvin, P. R. The renaissance of fluorescence resonance energy transfer. Nature structural biology 7 (9), 730–734 (2000).

[28] Gayrard, C. & Borghi, N. Fret-based molecular tension microscopy. Methods 94, 33–42 (2016).

[29] Colom, A. et al. A fluorescent membrane tension probe. Nature chemistry 10 (11), 1118–1125 (2018).

[30] Roffay, C., Chan, C. J., Guirao, B., Hiiragi, T. & Graner, F. Inferring cell junction tension and pressure from cell geometry. Development 148 (18), dev192773 (2021).

[31] Ishihara, S. et al. Comparative study of noninvasive force and stress inference methods in tissue. The European Physical Journal E 36 (4), 1–13 (2013).

[32] Thompson, D. W. & Thompson, D. W. On growth and form Vol. 2 (Cambridge university press Cambridge, 1942).

[33] Lecuit, T. & Lenne, P.-F. Cell surface mechanics and the control of cell shape, tissue patterns and morphogenesis. Nature reviews Molecular cell biology 8 (8), 633–644 (2007).

[34] Ishihara, S. & Sugimura, K. Bayesian inference of force dynamics during morphogenesis. Journal of theoretical biology 313, 201–211 (2012).

[35] Chiou, K. K., Hufnagel, L. & Shraiman, B. I. Mechanical stress inference for two dimensional cell arrays. PLoS computational biology 8 (5), e1002512 (2012).

[36] Nagai, T. & Honda, H. A dynamic cell model for the formation of epithelial tissues. Philosophical Magazine B 81 (7), 699–719 (2001)

[37] Farhadifar, R., Röper, J.-C., Aigouy, B., Eaton, S. & Jülicher, F. The influence of cell mechanics, cell-cell interactions, and proliferation on epithelial packing. Current Biology 17 (24), 2095–2104 (2007).

[38] Brodland, G. W. et al. Cellfit: a cellular force-inference toolkit using curvilinear cell boundaries. PloS one 9 (6), e99116 (2014).

[39] Kong, W. et al. Experimental validation of force inference in epithelia from cell to tissue scale. Scientific reports 9 (1), 1–12 (2019).

[40] Veldhuis, J. H. et al. Inferring cellular forces from image stacks. Philosophical Transactions of the Royal Society B: Biological Sciences 372 (1720), 20160261 (2017).

[41] Xu, M., Wu, Y., Shroff, H., Wu, M. & Mani, M. A scheme for 3-dimensional morphological reconstruction and force inference in the early c. elegans embryo. PLoS One 13 (7), e0199151 (2018).

[42] Noll, N., Streichan, S. J. & Shraiman, B. I. Variational method for image-based inference of internal stress in epithelial tissues. Physical Review X 10 (1), 011072 (2020).

[43] Boissonnat, J.-D. & Karavelas, M. I. On the combinatorial complexity of euclidean voronoi cells and convex hulls of d-dimensional spheres, Vol. 3, 305–312 (2003).

[44] Boissonnat, J.-D., Wormser, C. & Yvinec, M. in Curved voronoi diagrams 67–116 (Springer, 2006).

[45] Eppstein, D. A möbius-invariant power diagram and its applications to soap bubbles and planar lombardi drawing. Discrete & Computational Geometry 52 (3), 515–550 (2014)

[46] Sullivan, J. Nonspherical bubble clusters, 453–456 (2014).

[47] Liu, S., Lemaire, P., Munro, E. & Mani, M. A mathematical theory for the mechanics of three-dimensional cellular aggregates reveals the mechanical atlas for ascidian embryogenesis. bioRxiv 2022–11 (2022).

[48] Ichbiah, Sacha and Delbary, Fabrice and Turlier, Hervé. foambryo: 3d tension and pressure inference from microscopy images. URL https://github.com/VirtualEmbryo/foambryo.

[49] Batchelor, G. The stress system in a suspension of force-free particles. Journal of fluid mechanics 41 (3), 545–570 (1970).

[50] Meyer, M., Desbrun, M., Schröder, P. & Barr, A. H. in Discrete differential-geometry operators for triangulated 2-manifolds 35–57 (Springer, 2003).

[51] Crane, K. Discrete differential geometry: An applied introduction. Notices of the AMS, Communication 1153–1159 (2018).

[52] Brakke, K. A. The surface evolver. Experimental mathematics 1 (2), 141–165 (1992)

[53] Maître, J.-L. et al. Asymmetric division of contractile domains couples cell positioning and fate specification. Nature 536 (7616), 344–348 (2016).

[54] da Rocha, H. B., Bleyer, J. & Turlier, H. A viscous active shell theory of the cell cortex. Journal of the Mechanics and Physics of Solids 164, 104876 (2022).

[55] Da, F., Batty, C. & Grinspun, E. Multimaterial mesh-based surface tracking. ACM Trans. Graph. 33 (4), 112–1 (2014).

[56] Lorensen, W. E. & Cline, H. E. Marching cubes: A high resolution 3d surface construction algorithm. ACM siggraph computer graphics 21 (4), 163–169 (1987).

[57] Alliez, P. et al. in 3D mesh generation 5.5.1 edn (CGAL Editorial Board, 2022). URL https://doc.cgal.org/5.5.1/Manual/packages.html#PkgMesh3.

[58] Stringer, C., Wang, T., Michaelos, M. & Pachitariu, M. Cellpose: a generalist algorithm for cellular segmentation. Nature methods 18 (1), 100–106 (2021).

[59] Danielsson, P.-E. Euclidean distance mapping. Computer Graphics and image processing 14 (3), 227–248 (1980).

[60] Bai, M. & Urtasun, R. Deep watershed transform for instance segmentation, 5221–5229 (2017).

[61] Wang, W. et al. Learn to segment single cells with deep distance estimator and deep cell detector. Computers in biology and medicine 108, 133–141 (2019).

[62] Cousty, J., Bertrand, G., Najman, L. & Couprie, M. Watershed cuts: Minimum spanning forests and the drop of water principle. IEEE transactions on pattern analysis and machine intelligence 31 (8), 1362–1374 (2008).

[63] Kappes, J. H., Speth, M., Andres, B., Reinelt, G. & Schn, C. Globally optimal image partitioning by multicuts, 31–44 (Springer, 2011).

[64] Bailoni, A. et al. Gasp, a generalized framework for agglomerative clustering of signed graphs and its application to instance segmentation, 11645–11655 (2022).

[65] Wolf, S. et al. The mutex watershed: efficient, parameter-free image partitioning, 546–562 (2018).

[66] Ichbiah, S. & Turlier, H. Simulation dataset to benchmark 3D force inference methods (2023). URL https://doi.org/10.5281/zenodo.7881017.

[67] Ichbiah, Sacha and Turlier, Hervé. delaunay-watershed: multimaterial surface mesh reconstruction from 3d segmentation masks. URL https://github.com/VirtualEmbryo/delaunay-watershed.

[68] Harmand, N. Pertinence et limites des tensions de surface et de ligne pour rendre compte des formes des cellules épithéliales. Ph.D. thesis, Université de Paris (2019).

[69] Nocedal, J. & Wright, S. J. Numerical optimization (Springer, 1999).

[70] Dumollard, R., Hebras, C., Besnardeau, L. & McDougall, A. Beta-catenin patterns the cell cycle during maternal-to-zygotic transition in urochordate embryos. Dev Biol 384 (2), 331–342 (2013).

[71] Stewart, M. P. et al. Hydrostatic pressure and the actomyosin cortex drive mitotic cell rounding. Nature 469 (7329), 226–230 (2011)

[72] Taubenberger, A. V., Baum, B. & Matthews, H. K. The mechanics of mitotic cell rounding. Frontiers in cell and developmental biology 687 (2020).

[73] Graner, F. & Riveline, D. ‘the forms of tissues, or cell-aggregates’: D’arcy thompson’s influence and its limits. Development 144 (23), 4226–4237 (2017).

[74] Lemaire, P. Unfolding a chordate developmental program, one cell at a time: invariant cell lineages, short-range inductions and evolutionary plasticity in ascidians. Dev Biol 332 (1), 48–60 (2009).

[75] Sherrard, K., Robin, F., Lemaire, P. & Munro, E. Sequential activation of apical and basolateral contractility drives ascidian endoderm invagination. Current Biology 20 (17), 1499–1510 (2010).

[76] Hashimoto, H., Robin, F. B., Sherrard, K. M. & Munro, E. M. Sequential contraction and exchange of apical junctions drives zippering and neural tube closure in a simple chordate. Developmental cell 32 (2), 241–255 (2015).

[77] Hashimoto, H. & Munro, E. Differential expression of a classic cadherin directs tissue-level contractile asymmetry during neural tube closure. Developmental cell 51 (2), 158–172 (2019).

[78] Yamamoto, K. & Kimura, A. An asymmetric attraction model for the diversity and robustness of cell arrangement in nematodes. Development 144 (23), 4437–4449 (2017).

[79] Yamamoto, K. et al. Dissecting the subcellular forces sculpting early c. elegans embryos. bioRxiv 2023–03 (2023).

[80] Fernandez, R. et al. Imaging plant growth in 4d: robust tissue reconstruction and lineaging at cell resolution. Nature methods 7 (7), 547–553 (2010).

[81] Wolny, A. et al. Accurate and versatile 3d segmentation of plant tissues at cellular resolution. Elife 9, e57613 (2020).

[82] Kirillov, A. et al. Segment anything (2023). 2304.02643.

[83] Ebrahim, S. et al. Nmii forms a contractile transcellular sarcomeric network to regulate apical cell junctions and tissue geometry. Current biology 23 (8), 731–736 (2013).

[84] Zhu, M. et al. Developmental clock and mechanism of de novo polarization of the mouse embryo. Science 370 (6522), eabd2703 (2020)

[85] Turlier, H., Audoly, B., Prost, J. & Joanny, J.-F. Furrow constriction in animal cell cytokinesis. Biophysical journal 106 (1), 114–123 (2014).

[86] Hannezo, E., Prost, J. & Joanny, J.-F. Theory of epithelial sheet morphology in three dimensions. Proceedings of the National Academy of Sciences 111 (1), 27–32 (2014).

[87] Mayer, M., Depken, M., Bois, J. S., Jülicher, F. & Grill, S. W. Anisotropies in cortical tension reveal the physical basis of polarizing cortical flows. Nature 467 (7315), 617–621 (2010).

[88] Naganathan, S. R., Fürthauer, S., Nishikawa, M., Jülicher, F. & Grill, S. W. Active torque generation by the actomyosin cell cortex drives left–right symmetry breaking. Elife 3, e04165 (2014).

[89] Salbreux, G. & Jülicher, F. Mechanics of active surfaces. Physical Review E 96 (3), 032404 (2017).

[90] Torres-Sánchez, A., Millán, D. & Arroyo, M. Modelling fluid deformable surfaces with an emphasis on biological interfaces. Journal of fluid mechanics 872, 218–271 (2019).

[91] Ichbiah, S., Delbary, F. & Turlier, H. Differentiable rendering for 3d fluorescence microscopy. arXiv preprint arXiv:2303.10440 (2023).

[92] Sugimura, K., Lenne, P.-F. & Graner, F. Measuring forces and stresses in situ in living tissues. Development 143 (2), 186–196 (2016).

[93] Roca-Cusachs, P., Conte, V. & Trepat, X. Quantifying forces in cell biology. Nature cell biology 19 (7), 742–751 (2017).

[94] Munjal, A., Philippe, J.-M., Munro, E. & Lecuit, T. A self-organized biomechanical network drives shape changes during tissue morphogenesis. Nature 524 (7565), 351–355 (2015).

[95] Hannezo, E. & Heisenberg, C.-P. Mechanochemical feedback loops in development and disease. Cell 178 (1), 12–25 (2019).

[96] Bailles, A., Gehrels, E. W. & Lecuit, T. Mechanochemical principles of spatial and temporal patterns in cells and tissues. Annual review of cell and developmental biology 38, 321–347 (2022).

[97] McDougall, A., Lee, K. W. & Dumollard, R. Microinjection and 4d fluorescence imaging in the eggs and embryos of the ascidian phallusia mammillata. Methods Mol Biol 1128, 175–185 (2014).

[98] McDougall, A. et al. in Centrosomes and spindles in ascidian embryos and eggs, Vol. 129 317–339 (Elsevier, 2015).

