## Supplementary Note for "Embryo mechanics cartography: inference of 3D force atlases from fluorescence microscopy"

In this Supplementary Note, we provide additional theoretical and implementation details on the Delaunay-watershed mesh generation algorithm, the 3D active foam simulations, and the force inference pipeline.

#### A Multimaterial mesh generation

Our multimaterial nonmanifold triangle mesh data structure for surfaces is quite specific, and there is currently no library available to generate them directly. To overcome this difficulty, we used procedures to generate multimaterial tetrahedral volume meshes and developed methods to extract their boundaries as multimaterial triangle surface meshes.

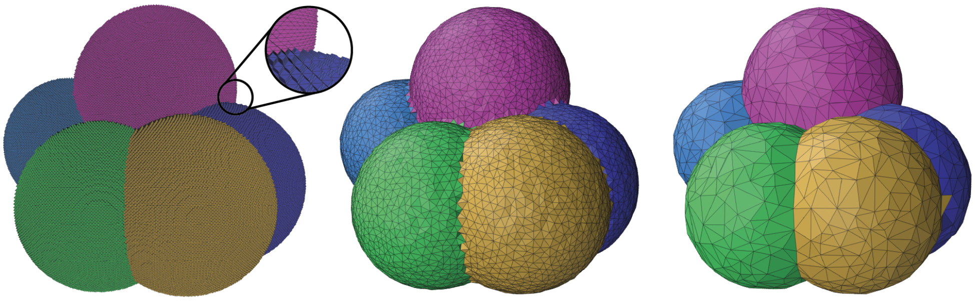

Instance segmentation CGAL Delaunay-Watershed

##### A.1 Instance segmentation

If we decompose each voxel into four tetrahedra, an instance segmentation can be viewed as a tetrahedral mesh. The task of generating a mesh from a voxel segmentation can thus be viewed as a remeshing procedure, where we convert a very fine mesh into a coarser one, which is better suited to our analysis.

##### A.2 CGAL-based 3D surface mesh generation

CGAL [1] provides a popular tetrahedral mesh generation algorithm, which has been used in biomechanical modeling, to extract volumetric meshes from 3D scans from different imaging modalities. It constitutes our baseline, from which we compared the results obtained with our proposed mesh generation algorithm, Delaunay-watershed.

##### A.3 Delaunay-watershed mesh generation algorithm (2D and 3D)

###### A.3.1 Distance transform map

Given an instance segmentation, we find the boundaries of the different instances with the function *find_boundaries* of the Python package scikit-image [11]. These borders can be converted into a distance map using a Euclidean distance transform with the function *distance_transform_edt* of the Python package *SciPy* [12].

##### A.3.2 Sampling of control points

To compute a tessellation of the space, we need to choose control points that will form the vertices of the tessellation. The points of interest lie into the borders of the objects, i.e. where the EDT is minimal. To regularize the tessellation, one can also add the local maxima. These extrema are efficiently computed using the *maxpool3d* function of *PyTorch* [9], with a kernel size of (d,d,d), $d \in \mathbb{Z}^+$, that will define the refinement of our mesh, and a stride of (1,1,1).

##### A.3.3 Delaunay tesselation

From these points, n-dimensional Delaunay tessellations are generated using the Delaunay function from the Python library scipy.spatial [12].

##### A.3.4 Dual Voronoi graph

The dual Voronoi graph $\mathcal{G} = (\mathcal{N}, \mathcal{L}, \mathcal{W})$ is then constructed from the tessellation, by looking at the adjacency of the simplices obtained. In this dual representation:

- In 2D, $\mathcal{N}$ contains triangles, $\mathcal{L}$ edges, and weights are calculated by averaging the value of the Euclidean distance transform of $n_s = 10$ points distributed regularly along the edge.
- In 3D, $\mathcal{N}$ contains the tetrahedra, $\mathcal{L}$ the triangles, and the weights are calculated by computed the average value of the euclidean distance transform of $n_s = 9$ regularly sampled points distributed within the triangle. This graph is converted into a NetworkX [5] graph, better suited to do operations such as node clustering.

##### A.3.5 Watershed algorithm

We use the algorithm 2 from [3] to implement the seeded watershed. The seeds are computed using the EDT: for each label $l$, we take the pixel $px_l^*$ corresponding to the global minimum of the EDT of the pixels having this label. However, to apply our watershed algorithm, the seed needs to be a node of our graph, i.e., a simplex of the tessellation: a tetrahedra in 3D or a triangle in 2D. For each label, we take the simplex of centroid c that minimizes the distance to the pixel $px_l^*$ previously found as a seed.

#### A.4 Discrete expressions of geometrical quantities on triangular surface meshes

Below, we provide analytical formulas for calculating several discrete geometrical quantities and their derivatives on triangular surface meshes. In the following, we will use Roman indices $(i, j, k)$ for quantities associated with cells (or regions), such as a tension $\gamma_{ij}$ between cells of indices $i$ and $j$, and Greek indices $\alpha$ for quantities associated with the mesh, such as the vertices denoted $\{\vec{x}_\alpha\}_{\alpha=1}^{n_v}$.

##### A.4.1 Interface area and derivatives

Each interface area $m$ and its gradient may be decomposed into a sum on its triangles

$$\mathcal{A}_m = \sum_{t \in \mathcal{A}_m} A_t, \qquad \frac{\partial \mathcal{A}_m}{\partial \vec{x}_\alpha} = \sum_{t \in \mathcal{A}_m} \frac{\partial A_t}{\partial \vec{x}_\alpha}. \tag{1}$$

Therefore, we provide the formula for only a single triangle, defined by its three vertices: $t = (\vec{x}_\alpha, \vec{x}_\beta, \vec{x}_\gamma)$. The non-unit and unit normals to the triangle $t$ are defined as

$$\vec{N}_t = (\vec{x}_\beta - \vec{x}_\alpha) \times (\vec{x}_\gamma - \vec{x}_\alpha), \qquad \vec{n}_t = \frac{\vec{N}_t}{\|\vec{N}_t\|}. \tag{2}$$

The area of the triangle $t$ follows as:

$$A_t = \frac{1}{2}\|\vec{N}_t\|. \tag{3}$$

To obtain the area gradient with respect to $\vec{x}_\alpha$, we calculate the variation of $A_t^2 = \frac{1}{4}\vec{N}_t \cdot \vec{N}_t$:

$$\delta\big(S_t^2\big) = \frac{1}{2}\vec{N}_t \cdot \delta\vec{N}_t = \frac{1}{2}\vec{N}_t \cdot \Big[(\vec{x}_\gamma - \vec{x}_\beta) \times \delta\vec{x}_\alpha\Big] = \frac{1}{2}\vec{N}_t \times \Big[(\vec{x}_\gamma - \vec{x}_\beta)\Big] \cdot \delta\vec{x}_\alpha.$$

From $\delta\big(A_t^2\big) = 2A_t\,\delta(A_t)$, we deduce

$$\frac{\partial A_t}{\partial \vec{x}_\alpha} = \frac{1}{2}\vec{n}_t \times (\vec{x}_\gamma - \vec{x}_\beta). \tag{4}$$

##### A.4.2 Cell volumes and derivatives

Stokes' theorem tells us that for a closed manifold, for any vectorial function f we have :

$$\iiint_{\mathcal{V}} \mathrm{div}(\vec{f})\mathrm{d}\tau = \oint_S \vec{f} \cdot \vec{n}\,\mathrm{d}s.$$

If we take $\vec{f}(\vec{x}) = \frac{1}{3}\vec{x}$, for which $div(\vec{f})(\vec{x}) = \frac{1}{3}(1+1+1)$, we have :

$$\mathcal{V} = \iiint_{\mathcal{V}} \mathrm{d}\tau = \frac{1}{3}\oint_S \vec{x} \cdot \vec{n}\,\mathrm{d}s = \frac{1}{3}\sum_{t=(\vec{x}_\alpha,\vec{x}_\beta,\vec{x}_\gamma)\in S} \iint_t \vec{x} \cdot \vec{n}\,\mathrm{d}s.$$

We will evaluate this integral on every triangle $t = (\vec{x}_\alpha, \vec{x}_\beta, \vec{x}_\gamma) \in S$.
The center of gravity of the triangle, defined as $\iint_{\vec{x}\in t}(\vec{x} - \vec{x}_G) \cdot \vec{n}\,\mathrm{d}s = 0$ reads

$$\vec{x}_G = \frac{\vec{x}_\alpha + \vec{x}_\beta + \vec{x}_\gamma}{3}. \tag{5}$$

We have

$$\iint_{\vec{x}\in t} \vec{x} \cdot \vec{n}\,\mathrm{d}s = \iint_{\vec{x}\in t} \vec{x}_G \cdot \vec{n}\,\mathrm{d}s = (\vec{n} \cdot \vec{x}_G)S_t = (\vec{n} \cdot \vec{x}_G)\frac{\|(\vec{x}_\gamma - \vec{x}_\alpha) \times (\vec{x}_\beta - \vec{x}_\alpha)\|}{2}.$$

By developing the expressions of $\vec{x}_G$ and $S_t$, we can write:

$$\begin{aligned}
\iint_{\vec{x}\in t} \vec{x} \cdot \vec{n}\,\mathrm{d}s &= \frac{1}{6}(\vec{x}_\alpha + \vec{x}_\beta + \vec{x}_\gamma) \cdot [(\vec{x}_\gamma - \vec{x}_\alpha) \times (\vec{x}_\beta - \vec{x}_\alpha)] \\
&= \frac{1}{6}(\vec{x}_\gamma - \vec{x}_\alpha + \vec{x}_\beta - \vec{x}_\alpha + 3\vec{x}_\alpha + \vec{x}_\beta + \vec{x}_\gamma) \cdot [(\vec{x}_\gamma - \vec{x}_\alpha) \times (\vec{x}_\beta - \vec{x}_\alpha)] \\
&= \frac{1}{2}\vec{x}_\alpha \cdot (\vec{x}_\gamma \times \vec{x}_\beta) \\
&= \frac{1}{2}\det(\vec{x}_\alpha, \vec{x}_\beta, \vec{x}_\gamma).
\end{aligned}$$

Finally,

$$\mathcal{V} = \frac{1}{3}\oint_S \vec{x} \cdot \vec{n}\,\mathrm{d}s = \frac{1}{6}\sum_{(\vec{x}_\alpha,\vec{x}_\beta,\vec{x}_\gamma)\in S} \det(\vec{x}_\alpha, \vec{x}_\beta, \vec{x}_\gamma). \tag{6}$$

Thus, the derivative of the volume with respect to the vertex writes

$$\frac{\partial \mathcal{V}}{\partial \vec{x}_\alpha} = \frac{1}{6} \sum_{(\vec{x}_\alpha,\vec{x}_\beta,\vec{x}_\gamma)\in triangles} \vec{x}_\beta \times \vec{x}_\gamma. \tag{7}$$

##### A.4.3 Junction length and derivatives

The length of a discrete curve can be written :

$$\mathcal{L} = \sum_{(\vec{x_\alpha},\vec{x_\beta})\in edges} \|\vec{x_\alpha} - \vec{x_\beta}\|. \tag{8}$$

Consider an edge $(\vec{x_\alpha}, \vec{x_\beta})$ of length $\mathrm{L}(\vec{x_\alpha}, \vec{x_\beta}) = \|\vec{x_\alpha} - \vec{x_\beta}\|$. To compute its derivative, we place ourselves on a basis where $\vec{e}_1 = \frac{\vec{x_\alpha}-\vec{x_\beta}}{\|\vec{x_\alpha}-\vec{x_\beta}\|}$ and

$$\mathrm{L}(\vec{x_\alpha} + \vec{\delta}, \vec{x_\beta}) = \|\vec{x_\alpha} - \vec{x_\beta}\| + \vec{\delta}\cdot\vec{e}_1.$$

The derivative with respect to each vertex is deduced as follows

$$\frac{\partial \mathrm{L}}{\partial \vec{x_\alpha}} = \frac{\vec{x_\alpha} - \vec{x_\beta}}{\|\vec{x_\alpha} - \vec{x_\beta}\|}. \tag{9}$$

##### A.4.4 Contact angles

For each junction $(i, j, k)$ between three cells (or the exterior), each edge is connected to three triangles, making the interface between two materials: $t_{ij}, t_{ik}, t_{jk}$ of normals $\vec{n}_{ij}, \vec{n}_{ik}, \vec{n}_{jk}$ (See Ext Fig. 1a)

$$\alpha_{ab} = \arccos(\vec{n}_{ac}\cdot\vec{n}_{bc}) \text{ for any (a,b,c) permutation of (i,j,k).} \tag{10}$$

For force inference the junction angles that appears in Young-Dupré force balance variants are defined as the average of the dihedral angles computed on every edges belonging to the junction.

##### A.4.5 Mean curvature

For a smooth manifold, the mean curvature $\bar{\kappa}$ at a given point is, by definition, the divergence of the normal vector at this point. A possible differential geometry definition of the curvature normal $H\,\vec{n}$ is

$$H\vec{n} = \lim_{\mathrm{diam(A)}\to 0} \frac{\vec{\nabla} A}{AS},$$

where $A$ is a small area around the point where the curvature is evaluated and diam(S) its diameter.

A discrete geometry equivalent to the previous formula was derived in [4] and reads

$$-H_i\,\vec{n}_i = \frac{1}{4}\sum_{j\in N_1(i)} \big(\cot\alpha_{ij} + \cot\beta_{ij}\big)(\vec{x}_j - \vec{x}_i), \tag{11}$$

where $\alpha_j$ and $\beta_j$ are the two angles opposite to the edge in the two triangles that share the edge $(\vec{x}_i, \vec{x}_j)$ as shown in the Ext Fig. 1d, and $N_1(i)$ is the set of neighbors with one ring of the vertex $i$.

For force inference, the interface mean curvatures that appears in Laplace equations are defined as the average of the mean-curvature computed on every vertices belonging to the interface.

### B Mechanical equilibrium in a heterogeneous foam

#### B.1 Heterogeneous foam model hypotheses

We assume that the shape of the blastomeres in the early embryos of interest is primarily controlled by cortical tensions [7, 8]. These surface tensions originate from cortical contractility and are considered homogeneous and isotropic at each cell-cell interface. The tension of the plasma membrane is lower, generically by an order of magnitude [7] but it may be counted in the surface tensions if necessary. The direct negative contribution to the surface tension of adhesion molecules (cadherin) is also negligible in general [7], but may also be counted in interface tensions, as long as all effective interface tensions remain strictly positive. Therefore, an assembly of cells is supposed to be akin to a heterogeneous foam-like structure but where each interface may have a different surface tension, which is controlled actively by adjacent cells. The evolution of cell shapes is assumed to be quasistatic, which means that viscous dissipation is neglected. This hypothesis is well justified by comparing the typical timescale associated with viscous relaxation of the cortex, on the order of dozens of seconds [6, 10], and the timescale associated with typical morphogenetic events in early embryos, rather than of the order of dozens of minutes. Moreover, we assume that blastomeres conserve their volumes on typical morphogenetic timescales (between two divisions).

#### B.2 Foam-like 3D simulations

##### B.2.1 Lagrangian function and force on a vertex

Our numerical simulations consist of a constrained optimization of a surface energy defined on an initial nonmanifold triangular mesh. The surface energy and Lagrangian function for a set of $n_c$ cells are defined as follows:

$$\mathcal{E} = \sum_{m=1}^{n_m} \gamma_m \mathcal{A}_m, \tag{12}$$

$$\mathcal{L} = \mathcal{E} - \sum_{k=1}^{n_c} p_k (\mathcal{V}_k - \mathcal{V}_k^0), \tag{13}$$

where $\gamma_m$ and $\mathcal{A}_m$ are, respectively, the surface tension and area of the interface between the regions $a_m$ and $b_m$ where $a_m$, $b_m \in [0, n_c]$, $m \in [1, n_m]$ are an index of a membrane, which identifies a pair of regions. $p_k$ and $\mathcal{V}_k^0$ are, respectively, the pressure and target volume value of the cell $k$. Note that for interfaces $a_m$ and $b_m$ span $[0, n_c]$, where 0 refers to the external medium, while for cells $k$ span $[1, n_c]$.

From the Lagrangian function, one can calculate the force $\vec{f}_x = -\dfrac{\partial \mathcal{L}}{\partial \vec{x}}$ at each vertex of the mesh $\vec{x} \in \{\vec{x}_\alpha\}_{\alpha=1}^{n_v}$ as follows

$$\vec{f}_x = -\frac{\partial \mathcal{E}}{\partial \vec{x}} + \sum_{k=1}^{n_c} p_k \frac{\partial \mathcal{V}_k}{\partial \vec{x}} = -\sum_{m=1}^{n_m} \gamma_m \frac{\partial \mathcal{A}_m}{\partial \vec{x}} + \sum_{k=1}^{n_c} p_k \frac{\partial \mathcal{V}_k}{\partial \vec{x}}. \tag{14}$$

##### B.2.2 Lagrangian function for embryos within a shell

For the example of C. Elegans, the optimization of a surface energy must satisfy an additional constraint, that is, that every vertex must lie in the shell. We call $\mathcal{S}$ the shell, $\kappa$ the stiffness of the shell, $\mathcal{F}_{el}$ the elastic force exerted by the shell on the vertices outside its bounds, defined by :

$$\mathcal{F}_{el}(\vec{x}_\alpha) = \|\vec{x}_\alpha - \vec{x}_{\mathcal{S}}(\vec{x}_\alpha)\|^2 \cdot \mathbf{N}_{\mathcal{S}}(\vec{x}_{\mathcal{S}}(\vec{x}_\alpha)), \tag{15}$$

where $\mathbf{N}_{\mathcal{S}}(\vec{x})$ is the normal of the surface of the shell at the point $\vec{x}$ (belonging to the shell), and $\vec{x}_{\mathcal{S}}(\vec{x})$ is the orthogonal projection of the vertex $\vec{x}$ on the surface of the shell $\mathcal{S}$.

The surface energy and Lagrangian function for a set of $N$ cells then take the following form

$$\mathcal{E} = \sum_{m=1}^{n_m} \gamma_m \mathcal{A}_m + \frac{\kappa}{2} \sum_{\tilde{x}_\alpha \notin \mathcal{S}} \mathcal{F}_{el}(\vec{x}_\alpha), \tag{16}$$

$$\mathcal{L} = \mathcal{E} - \sum_{k=1}^{n_c} p_k (\mathcal{V}_k - \mathcal{V}_k^0). \tag{17}$$

#### B.3 Variants of the Young-Dupré tension balance

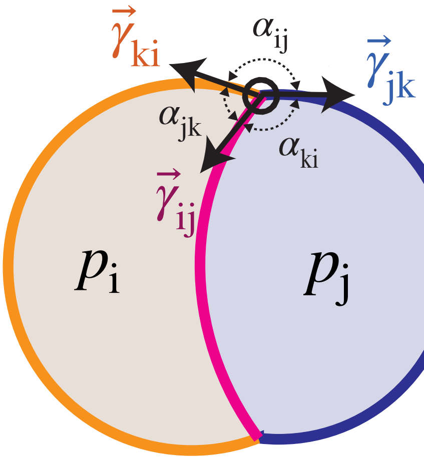


To obtain scalar values of the tensions, we first choose to fix the average surface tension to 1, with a first equation:

$$\sum_{m=1}^{n_m} \gamma_m = n_m. \tag{18}$$

Tension-balance gives relative relationships between surface tensions, and can be expressed in numerous ways. For all the following formulas, $(i, j, k)$ represents three materials (cells or exterior media) that meet at a triple junction.

***Young-Dupré***

$$\forall (i,j,k), \quad \gamma_{ij} + \gamma_{ik} \cos \alpha_{jk} + \gamma_{jk} \cos \alpha_{ik} = 0 \tag{19}$$

$$\forall (i,j,k), \quad \gamma_{ij} \cos \alpha_{jk} + \gamma_{ik} + \gamma_{jk} \cos \alpha_{ij} = 0 \tag{20}$$

$$\forall (i,j,k), \quad \gamma_{ij} \cos \alpha_{ik} + \gamma_{ik} \cos \alpha_{ij} + \gamma_{jk} = 0 \tag{21}$$

***Projection Young-Dupré***

$$\forall (i,j,k), \quad \gamma_{ij} + \gamma_{ik} \cos \alpha_{jk} + \gamma_{jk} \cos \alpha_{ik} = 0 \tag{22}$$

$$\forall (i,j,k), \quad \gamma_{ik} \sin \alpha_{jk} - \gamma_{jk} \sin \alpha_{ik} = 0 \tag{23}$$

***Lami***

$$\forall(i,j,k), \quad \frac{\gamma_{ij}}{\sin\alpha_{ij}} = \frac{\gamma_{ik}}{\sin\alpha_{ik}} = \frac{\gamma_{jk}}{\sin\alpha_{jk}} \tag{24}$$

***Inverse Lami***

$$\forall(i,j,k), \quad \gamma_{ij}\sin\alpha_{ik} = \gamma_{ik}\sin\alpha_{ij} \tag{25}$$

$$\forall(i,j,k), \quad \gamma_{ij}\sin\alpha_{jk} = \gamma_{jk}\sin\alpha_{ij} \tag{26}$$

$$\forall(i,j,k), \quad \gamma_{ik}\sin\alpha_{jk} = \gamma_{jk}\sin\alpha_{ik} \tag{27}$$

***Lami logarithm***

$$\forall(i,j,k), \log\gamma_{ij} - \log\sin\alpha_{ij} = \log\gamma_{ik} - \log\sin\alpha_{ik} = \log\gamma_{jk} - \log\sin\alpha_{jk} \tag{28}$$

#### B.4 Mesh-based variational force balance

In this study, we propose new force-balance equations, *Variational Young-Dupré* and *Variational Laplace* equations, that can be used for tension inference. In this section we propose a detailed derivation of these equations. Our mesh, which describes the surface of the embryo, is composed of $n_v$ vertices, $n_c$ cells of volumes $V_l^0$ for $l \in 1 \ldots n_c$ and of $n_m$ membranes (or interfaces) of area $A_m$. The equilibrium shape is given by minimizing the energy of the membranes, which have a defined surface tension $\gamma_m$, under the constraint of volume conservation $\mathcal{V}$. We call the set of vertices of the mesh $\mathbb{V} = \{\vec{x}_\alpha\}_{\alpha=1}^{n_v}$,the set of cells $\mathcal{C}$, and the set of membranes $\mathcal{M}$.

##### B.4.1 Geometrical model

Keeping the same notation as before, the Lagrangian function that is minimized is:

$$\mathcal{L} = \sum_{m=1}^{n_m} \gamma_m \mathcal{A}_m - \sum_{k=1}^{n_c} p_k(\mathcal{V}_k - \mathcal{V}_k^0), \tag{29}$$

In equilibrium, we have $\dfrac{\partial\mathcal{L}}{\partial\vec{x}_\alpha} = \vec{0}$.

The equilibrium of the forces gives us, for each vertex $\vec{x}_\alpha$, the following equation :

$$\vec{0} = \sum_{m=1}^{n_m} \gamma_m \frac{\partial\mathcal{A}_m}{\partial\vec{x}} - \sum_{k=1}^{n_c} p_k \frac{\partial\mathcal{V}_k}{\partial\vec{x}}. \tag{30}$$

This can be summarized as a product between a huge tensor $Q \in \mathbb{R}^{n_v \times (n_m+n_c)\times 3}$ that contains the derivatives of the areas and volumes with respect to each vertex, and the matrix $\Pi \in \mathbb{R}^{n_m+n_c}$ that contains the surface tensions and pressures. This product gives the equilibrium equations for each of the vertices. The tensor Q can be conveniently expressed as a matrix of $\mathbb{R}^3$ vectors:

$$Q \times \Pi = \begin{pmatrix} \nabla_{\vec{x_1}}\mathcal{A}_1 & \ldots & \nabla_{\vec{x_1}}\mathcal{A}_{n_m} & \nabla_{\vec{x_1}}\mathcal{V}_1 & \ldots & \nabla_{\vec{x_1}}\mathcal{V}_{n_c} \\ . & & . & . & & . \\ . & & . & . & & . \\ . & & . & . & & . \\ . & & . & . & & . \\ \nabla_{\vec{x_{n_v}}}\mathcal{A}_1 & \ldots & \nabla_{\vec{x_{n_v}}}\mathcal{A}_{n_m} & \nabla_{\vec{x_{n_v}}}\mathcal{V}_1 & \ldots & \nabla_{\vec{x_{n_v}}}\mathcal{V}_{n_c} \end{pmatrix} \begin{pmatrix} \gamma_1 \\ \ldots \\ \gamma_{n_m} \\ -p_1 \\ \ldots \\ -p_{n_c} \end{pmatrix} = 0$$

##### B.4.2 State Equations

We will define a matrix $R \in \mathbb{R}^{(n_c+n_m)\times(n_c+n_m)} : R = Q^T Q$. We note that $R \times \Pi = 0$.

$$R = \begin{pmatrix} \sum_{i\in\mathbb{V}} \|\nabla_{\vec{x_i}}\mathcal{A}_1\|^2 & \dots & \sum_{i\in\mathbb{V}}(\nabla_{\vec{x_i}}\mathcal{A}_1 \cdot \nabla_{\vec{x_i}}\mathcal{A}_{n_m}) & \sum_{i\in\mathbb{V}}(\nabla_{\vec{x_i}}\mathcal{A}_1 \cdot \nabla_{\vec{x_i}}\mathcal{V}_1) & \dots & \sum_{i\in\mathbb{V}}(\nabla_{\vec{x_i}}\mathcal{A}_1 \cdot \nabla_{\vec{x_i}}\mathcal{V}_{n_c}) \\ \vdots & & \vdots & \vdots & & \vdots \\ \sum_{i\in\mathbb{V}}(\nabla_{\vec{x_i}}\mathcal{A}_{n_m} \cdot \nabla_{\vec{x_i}}\mathcal{A}_1) & \dots & \sum_{i\in\mathbb{V}} \|\nabla_{\vec{x_i}}\mathcal{A}_{n_m}\|^2 & \sum_{i\in\mathbb{V}}(\nabla_{\vec{x_i}}\mathcal{A}_{n_m} \cdot \nabla_{\vec{x_i}}\mathcal{V}_1) & \dots & \sum_{i\in\mathbb{V}}(\nabla_{\vec{x_i}}\mathcal{A}_{n_m} \cdot \nabla_{\vec{x_i}}\mathcal{V}_{n_c}) \\ \sum_{i\in\mathbb{V}}(\nabla_{\vec{x_i}}\mathcal{V}_1 \cdot \nabla_{\vec{x_i}}\mathcal{A}_1) & \dots & \sum_{i\in\mathbb{V}}(\nabla_{\vec{x_i}}\mathcal{V}_1 \cdot \nabla_{\vec{x_i}}\mathcal{V}_1) & \sum_{i\in\mathbb{V}} \|\nabla_{\vec{x_i}}\mathcal{V}_1\|^2 & \dots & \sum_{i\in\mathbb{V}}(\nabla_{\vec{x_i}}\mathcal{V}_1 \cdot \nabla_{\vec{x_i}}\mathcal{V}_{n_c}) \\ \vdots & & \vdots & \vdots & & \vdots \\ \sum_{i\in\mathbb{V}}(\nabla_{\vec{x_i}}\mathcal{V}_{n_c} \cdot \nabla_{\vec{x_i}}\mathcal{A}_1) & \dots & \sum_{i\in\mathbb{V}}(\nabla_{\vec{x_i}}\mathcal{V}_{n_c} \cdot \nabla_{\vec{x_i}}\mathcal{A}_{n_m}) & \sum_{i\in\mathbb{V}}(\nabla_{\vec{x_i}}\mathcal{V}_{n_c} \cdot \nabla_{\vec{x_i}}\mathcal{V}_1) & \dots & \sum_{i\in\mathbb{V}} \|\nabla_{\vec{x_i}}\mathcal{V}_{n_c}\|^2 \end{pmatrix}$$

We note that matrix R has a kernel of dimension 1. We need to add another equation to fix the tension scales (by setting the mean to 1 for example).

The state equations that we choose are $\mathcal{S} = R \times \Pi = 0$ which gives us discretized equations:

$$\forall l \in \mathcal{M}, \quad 0 = \sum_{m=1}^{n_m} \gamma_m \left( \sum_{i\in\mathbb{V}} \frac{\partial \mathcal{A}_m}{\partial \vec{x}_i} \cdot \frac{\partial \mathcal{A}_l}{\partial \vec{x}_i} \right) - \sum_{k=1}^{n_c} p_n \left( \sum_{i\in\mathbb{V}} \frac{\partial \mathcal{V}_k}{\partial \vec{x}_i} \cdot \frac{\partial \mathcal{A}_l}{\partial \vec{x}_i} \right) \tag{31}$$

$$\forall n \in \mathcal{C}, \quad 0 = \sum_{m=1}^{n_m} \gamma_m \left( \sum_{i\in\mathbb{V}} \frac{\partial \mathcal{A}_m}{\partial \vec{x}_i} \cdot \frac{\partial \mathcal{V}_n}{\partial \vec{x}_i} \right) - \sum_{k=1}^{n_c} p_n \left( \sum_{i\in\mathbb{V}} \frac{\partial \mathcal{V}_k}{\partial \vec{x}_i} \cdot \frac{\partial \mathcal{V}_n}{\partial \vec{x}_i} \right) \tag{32}$$

##### B.4.3 Matrix equations

We will note $\Gamma = (\gamma_1, \gamma_2 \dots \gamma_{n_m})^T$ the generalized vector of tensions and $\mathrm{P} = (p_1, p_2 \dots p_{n_c})^T$ the generalized vector of pressures.

We can represent the first set of discretized equations using matrix notation $\mathrm{G}_\Gamma \in \mathbb{R}^{n_m \times n_m}$, and $\mathrm{B}_P \in \mathbb{R}^{n_m \times n_c}$ verifying $\mathrm{G}_\Gamma \Gamma - \mathrm{B}_P \mathrm{P} = 0$ :

$$\forall l, m \in (1 \dots n_m) \quad \mathrm{G}_\Gamma^{lm} = \sum_{i\in\mathbb{V}} \frac{\partial \mathcal{A}_m}{\partial \vec{x}_i} \cdot \frac{\partial \mathcal{A}_l}{\partial \vec{x}_i}$$

$$\forall k \in (1 \dots n_c), l \in (1 \dots n_m) \quad \mathrm{B}_P^{lk} = \sum_{i\in\mathbb{V}} \frac{\partial \mathcal{V}_k}{\partial \vec{x}_i} \cdot \frac{\partial \mathcal{A}_l}{\partial \vec{x}_i}$$

$\mathrm{G}_\Gamma$ is generally not invertible; all tensions are fixed to one parameter.

We can represent the second set of discretized equations using matrix notation $\mathrm{B}_\Gamma \in \mathbb{R}^{n_c \times n_m}$, and $\mathrm{G}_P \in \mathbb{R}^{n_c \times n_c}$ verifying $\mathrm{B}_\Gamma \Gamma - \mathrm{G}_P \mathrm{P} = 0$ :

$$\forall n \in (1 \dots n_c), m \in (1 \dots n_m) \quad \mathrm{B}_\Gamma^{nm} = \sum_{i\in\mathbb{V}} \frac{\partial \mathcal{A}_m}{\partial \vec{x}_i} \cdot \frac{\partial \mathcal{V}_n}{\partial \vec{x}_i}$$

$$\forall k, n \in (1 \dots n_c) \quad \mathrm{G}_P^{nk} = \sum_{i\in\mathbb{V}} \frac{\partial \mathcal{V}_k}{\partial \vec{x}_i} \cdot \frac{\partial \mathcal{V}_n}{\partial \vec{x}_i}$$

$\mathrm{G}_P$ is invertible thus knowing $\Gamma$ and K, we can deduce P from these equations, that we call *Variational Laplace* equations:

$$\mathrm{P} = (\mathrm{G}_P)^{-1} \mathrm{B}_\Gamma \Gamma. \tag{33}$$

We can write $\Pi = \begin{pmatrix} \Gamma \\ \mathrm{P} \end{pmatrix}$ and write the previous equations into a block matrix M:

$$\mathrm{M} \times \Pi = 0 \iff \begin{pmatrix} \mathrm{G}_\Gamma & -\mathrm{B}_P \\ \mathrm{B}_\Gamma & -\mathrm{G}_P \end{pmatrix} \times \begin{pmatrix} \Gamma \\ \mathrm{P} \end{pmatrix} = 0 \tag{34}$$

Thus, the *Variational Young-Dupré* equations for tension inference are

$$(\mathrm{G}_\Gamma - \mathrm{B}_P(\mathrm{G}_P)^{-1}\mathrm{B}_\Gamma) \times \Gamma = 0, \tag{35}$$

$$(1 \cdots 1) \times \Gamma = n_m. \tag{36}$$

This last equation fixes the average of the surface tensions to 1.

##### B.4.4 Proof that the matrix $\mathbf{G}_P$ is invertible

In the previous section, we assumed that the matrix $\mathrm{G}^p$ is invertible. To demonstrate this hypothesis, we have to show that the matrix of volume gradients has full rank (linearly independent columns). For simplicity, assume that each face of each discretized cell has an interior point and that the cells are numbered in "decreasing" order, from the outermost(s) to the innermost(s), more precisely, in the sense that there exists $\vec{x}_{i_1} \in \mathrm{Cell}_1$ such that $\vec{x}_{i_1} \notin \bigcup_{k>=2} \mathrm{Cell}_k$, and recursively, for all $k \in (1 \ldots n_c)$, there exists $\vec{x}_{i_k} \in \mathrm{Cell}_k$ such that $\vec{x}_{i_k} \notin \bigcup_{k'>=k+1} \mathrm{Cell}_{k'}$. Now assume that the volume gradients are not linearly independent, then there exists $P_1, \ldots, P_{n_c} \in \mathbb{R}$ such that for all $i \in (1 \ldots n_v)$

$$\sum_{k=1}^{n_c} P_i \nabla_{\vec{x}_i} \mathcal{V}_k = 0. \tag{37}$$

Hence, for $i = i_1$, we simply have $P_1 \nabla_{\vec{x}_{i_1}} \mathcal{V}_1 = 0$. Furthermore, the gradient of $\mathcal{V}_1$ with respect to $\vec{x}_{i_1}$ is the sum of the gradients with respect to $\vec{x}_{i_1}$ of the volumes of the tetrahedra defined by the apex 0 and the triangles surrounding $\vec{x}_{i_1}$, so $\nabla_{\vec{x}_{i_1}} \mathcal{V}_1 \neq 0$ so that $P_1 = 0$ and

$$\sum_{k=2}^{n_c} P_i \nabla_{\vec{x}_i} \mathcal{V}_k = 0. \tag{38}$$

Thus, by recursion, we have $P_i = 0$ for all $i \in (1 \ldots n_c)$, so that the volume gradients are linearly independent. Consequently, the matrix $\mathrm{G}^p$ is invertible.

### C Force inference

#### C.1 Hypotheses for tension and pressure inference

We assume that each pressure $p$ is positive and put the pressure of the exterior medium at 0. The surface tensions $y$ are also positive, and as explained before we arbitrarily put their mean at 1. This choice of scale for the tensions imposes a scale on the pressures, as appears in the Laplace law $2\gamma_{ij} H_{ij} = p_i - p_j$. However, the scale of the mean curvature $H_{ij}$ depends on the size of the embryo. To compute the curvature with the right scaling, one would need to have a mesh that has the precise dimensions of the biological sample. This is not out of reach experimentally, but since we were interested in relative and not absolute values (absolute measurement of surface tensions being not available in any way), we found it more convenient to fix the biggest pressure at 1.

#### C.2 In silico validation

With our previously introduced 3D active foam model, we generated 47 simulations of cell clusters, ranging from 2 to 11 cells, with random tensions.

From these meshes, we generated artificial images by defining a bounding box and convolving the mesh by a Gaussian Point Spread Function. To do the convolution efficiently, we make the operation in Fourier space before doing an inverse Fourier transform to go back into the spatial domain.

A simple thresholding of these images allowed us to compute the membrane boundaries. The segmentation masks were then computed using a watershed algorithm. Masks were expanded until they merged to avoid holes inside embryos that would have perturbed meshing algorithms, and thus the computation of geometrical quantities.

#### C.3 Ordinary Least-Squares

Each of the linear problem encountered is in the following form

$$AX = C \tag{39}$$

where we want to determine the vector $X \in \mathbb{R}^m$ from $A \in \mathbb{R}^{n\times(m+1)}$, which is a rectangular matrix and the right-hand side $C \in \mathbb{R}^{m+1}$.

To determine an estimator of $X$, we choose the ordinary least-squares (OLS) method that minimizes the sum of squared residuals $\|AX - C\|^2$. The solution can be expressed in a closed form using the pseudo-inverse matrix $(A^TA)^{-1}A^T$:

$$X = (A^TA)^{-1}A^TC \tag{40}$$

In practice, we solve the ordinary least squares problem using the function linalg.lstq from the Python library SciPy [12].

#### C.4 Ordinary least-squares for tension inference

##### C.4.1 Form of the matrices A and C for tension inference

The form of the matrix $A$ and $C$ depends on the number of equations $n_{eq}$, which depends on the formula chosen.

$$A = \begin{pmatrix} \alpha_{11} & \dots & \alpha_{1n_m} \\ \dots & & \dots \\ \alpha_{n_{eq}1} & \dots & \alpha_{n_{eq}n_m} \\ 1 & \dots & 1 \end{pmatrix}, \quad X = \begin{pmatrix} \Gamma \end{pmatrix}, \quad C = \begin{pmatrix} 0 \\ \dots \\ 0 \\ n_m \end{pmatrix} \tag{41}$$

Note that A is very sparse, as each line has only three non-zero terms (except the last one).

##### C.4.2 Weighting by the junction length

For the tension inference equations (except for the variational Young-Dupré variant), each equation is linked to a given triple junction. The longer the junction, the better the determination of the angles of the junction will be. We decided to weight each of the equations by the length of the junction associated with it. Thus, if we consider the original unweighted matrices $A$ and $C$ used to infer tension, instead of minimizing $\|AX - C\|^2 = \sum_i((AX)_i - C_i)^2$ we minimize:

$$\sum_i ((AX)_i - C_i)^2 * L_i^2, \tag{42}$$

where $L_i$ is the length of the junction considered in the $i^{\text{th}}$ equation.

Here is a table of the median relative error of the different methods of tension inference, with and without weighting of junction length:

| | Young-Dupré | Proj. Young-Dupré | Lami | Inverse Lami | Lami Log |
|---|---|---|---|---|---|
| With length weight (%) | 10.1 | 10.4 | 11.7 | 11.0 | 11.8 |
| No length weight (%) | 10.8 | 11.7 | 12.7 | 11.9 | 12.4 |

##### C.4.3 Form with variational equations

With *Variational Young-Dupré* equations, we can express $A$ and $C$ as block matrices:

$$A = \begin{pmatrix} \mathrm{G}_\Gamma - \mathrm{B}_P(\mathrm{G}_P)^{-1}\mathrm{B}_\Gamma \\ 1 \end{pmatrix}, \quad X = \begin{pmatrix} \Gamma \end{pmatrix}, \quad C = \begin{pmatrix} 0 \\ n_m \end{pmatrix} \tag{43}$$

#### C.5 Ordinary Least Squares for pressure inference

##### C.5.1 Variational pressure inference

The *Variational Laplace* equations offers a way to infer pressures.

$$\mathrm{G}_P \mathrm{P} = \mathrm{B}_\Gamma \Gamma. \tag{44}$$

It is an ordinary least square problem with $A = \mathrm{G}_P, X = \mathrm{P}, C = \mathrm{B}_\Gamma \Gamma$.

##### C.5.2 Form of the matrices A and C for pressure inference with Laplace Law

There is one Laplace equation for each interface, so $n_m$ equations in total, thus $A \in \mathbb{R}^{n_m \times n_c}, X \in \mathbb{R}^{n_c}, C \in \mathbb{R}^{n_m}$. Each relation writes $p_{a_m} - p_{b_m} = \gamma_m H_m$ where $a_m$ and $b_m$ are the regions delimited by the interface $m$, of surface tension $\gamma_m$ and mean curvature $H_m$. Thus $A$ and $C$ writes:

$$A = \begin{pmatrix} \beta_{11} & \dots & \beta_{1n_c} \\ \dots & & \dots \\ \beta_{n_m 1} & \dots & \beta_{n_m n_c} \end{pmatrix}, \quad X = \begin{pmatrix} \mathrm{P} \end{pmatrix}, \quad C = \begin{pmatrix} 2\gamma_1 H_{a_1 b_1} \\ \dots \\ 2\gamma_{n_m} H_{a_{n_m} b_{n_m}} \end{pmatrix} \tag{45}$$

,

Where $\beta_{ij} = 1$ if $j \in \{a_i, b_i\}$ and 0 otherwise.

##### C.5.3 Weighting by the interfacial area

For the pressure inference equations with Laplace's law, each equation is linked to a given interface. The larger the area, the better the determination of the curvature of the interface will be. We decided to weight each of the equations by the area of the interface associated with it.

Thus, if we consider the original unweighted matrices $A$ and $C$ used to infer pressure, instead of minimizing $\|AX - C\|^2 = \sum_i ((AX)_i - C_i)^2$ we will minimize:

$$\sum_i ((AX)_i - C_i)^2 * \mathcal{A}_i^2, \tag{46}$$

where $\mathcal{A}_i$ is the area of the interface considered in the $i^{\text{th}}$ equation.

Here is a table of the median relative error of the different methods of pressure inference, with the tensions previously inferred using the Young-Dupré formula:

| | Laplace's Law | Variational Laplace |
|---|---|---|
| With area weight (%) | 12.5 | - |
| No area weight (%) | 12.6 | 12.0 |

### D Stress tensor

We consider a cell $a$. The cell surface is composed of membrane portion $m \in a$ of constant surface tension $\gamma_m^a$. The energy of the total foam writes $\mathcal{E} = \sum_{m=1}^{n_m} \gamma_m \mathcal{A}_m$, but alternatively, we can express the energy of the foam by summing up the contributions of each cell:

$$\mathcal{E} = \sum_a \sum_{m \in a} \gamma_m^a \mathcal{A}_m \tag{47}$$

For a membrane portion m that is in contact with the exterior medium, we can identify $\gamma_m$ and $\gamma_m^a$. However, for a surface shared between two cells, each membrane area is counted twice, so $\gamma_m^a = \frac{\gamma_m}{2}$.

We note $\mathcal{V}_{a+}$ the total volume of the cell and $\mathcal{V}_{a-}$ the volume of the cell enclosed by the membrane. We note $A_0$ the surface of the membrane The mean stress writes :

$$\overline{\underline{\underline{\sigma}}}_a = \frac{1}{\mathcal{V}_a} \int_{\mathcal{V}_{a+}} \underline{\underline{\sigma}}(\vec{x})\,\mathrm{d}V \tag{48}$$

We decompose it between the bulk and membrane components :

$$\overline{\underline{\underline{\sigma}}}_a = \frac{1}{\mathcal{V}_a} \int_{\mathcal{V}_{a-}} \underline{\underline{\sigma}}(\vec{x})\,\mathrm{d}V + \frac{1}{\mathcal{V}_a}\left( \int_{\mathcal{V}_{a+}} \underline{\underline{\sigma}}(\vec{x})\mathrm{d}V - \int_{\mathcal{V}_{a-}} \underline{\underline{\sigma}}(\vec{x})\mathrm{d}V \right) \tag{49}$$

The stress $\underline{\underline{\sigma}}$ is constant within the cell, its value being defined by hydrostatic pressure $p_a$. In the cell membrane, the small thickness limit established in [2] leads to the following relation:

$$\left( \int_{\mathcal{V}_{a+}} \underline{\underline{\sigma}}(\vec{x})\,\mathrm{d}V - \int_{\mathcal{V}_{a-}} \underline{\underline{\sigma}}(\vec{x})\,\mathrm{d}V \right) = \int_{A_0} \gamma^a (\underline{\underline{\delta}} - \vec{n} \otimes \vec{n})\,\mathrm{d}S \tag{50}$$

We note $\underline{\underline{\delta}}$ the identity tensor and $\vec{n} = \begin{bmatrix} n_1 \\ n_2 \\ n_3 \end{bmatrix}$ and $\vec{n} \otimes \vec{n} = \begin{bmatrix} n_1n_1 & n_1n_2 & n_1n_3 \\ n_2n_1 & n_2n_2 & n_2n_3 \\ n_3n_1 & n_3n_2 & n_3n_3 \end{bmatrix}$.

Eventually, the mean stress reads

$$\overline{\underline{\underline{\sigma}}}_a = -p_a \underline{\underline{\delta}} + \sum_{m \in a} \frac{\gamma_m^a}{\mathcal{V}_a} \int_m (\underline{\underline{\delta}} - \vec{n} \otimes \vec{n})\,\mathrm{d}S \tag{51}$$

We can decompose each surface membrane $m$ into individual triangles that make up it. As the normal is constant on a triangle, the integral simplifies itself :

$$\int_m (\underline{\underline{\delta}} - \vec{n} \otimes \vec{n})\,\mathrm{d}S = \sum_{t \in m} \int_t (\underline{\underline{\delta}} - \vec{n} \otimes \vec{n})\,\mathrm{d}S = \sum_{t \in m} A_t (\underline{\underline{\delta}} - \vec{n}_t \otimes \vec{n}_t) \tag{52}$$

$$\overline{\underline{\underline{\sigma}}}_a = -p_a \underline{\underline{\delta}} + \sum_{m \in a} \frac{\gamma_m^a}{\mathcal{V}_a} \sum_{t \in m} A_t (\underline{\underline{\delta}} - \vec{n}_t \otimes \vec{n}_t) \tag{53}$$

We can compute it numerically from our mesh once the force inference was performed and plot the tensor decomposed on its principal axes (eigenvectors) in each cell. Note that the eigenvalues may be positive, which corresponds to an extensile stress (in blue) or negative, which corresponds to a compressive stress (in red).

### References


[1] Pierre Alliez, Clément Jamin, Laurent Rineau, Stéphane Tayeb, Jane Tournois, and Mariette Yvinec. 3D mesh generation. In *CGAL User and Reference Manual*. CGAL Editorial Board, 5.5.1 edition, 2022.

[2] GK Batchelor. The stress system in a suspension of force-free particles. *Journal of fluid mechanics*, 41(3):545–570, 1970.

[3] Jean Cousty, Gilles Bertrand, Laurent Najman, and Michel Couprie. Watershed cuts: Minimum spanning forests and the drop of water principle. *IEEE transactions on pattern analysis and machine intelligence*, 31(8):1362–1374, 2008.

[4] Mathieu Desbrun, Mark Meyer, Peter Schröder, and Alan H Barr. Implicit fairing of irregular meshes using diffusion and curvature flow. In *Proceedings of the 26th annual conference on Computer graphics and interactive techniques*, pages 317–324, 1999.

[5] Aric Hagberg, Pieter Swart, and Daniel S Chult. Exploring network structure, dynamics, and function using networkx. Technical report, Los Alamos National Lab.(LANL), Los Alamos, NM (United States), 2008.

[6] Nargess Khalilgharibi, Jonathan Fouchard, Nina Asadipour, Ricardo Barrientos, Maria Duda, Alessandra Bonfanti, Amina Yonis, Andrew Harris, Payman Mosaffa, Yasuyuki Fujita, et al. Stress relaxation in epithelial monolayers is controlled by the actomyosin cortex. *Nature physics*, 15(8):839–847, 2019.

[7] Jean-Léon Maître, Ritsuya Niwayama, Hervé Turlier, François Nédélec, and Takashi Hiiragi. Pulsatile cell-autonomous contractility drives compaction in the mouse embryo. *Nature cell biology*, 17(7):849–855, 2015.

[8] Jean-Léon Maître, Hervé Turlier, Rukshala Illukkumbura, Björn Eismann, Ritsuya Niwayama, François Nédélec, and Takashi Hiiragi. Asymmetric division of contractile domains couples cell positioning and fate specification. *Nature*, 536(7616):344–348, 2016.

[9] Adam Paszke, Sam Gross, Francisco Massa, Adam Lerer, James Bradbury, Gregory Chanan, Trevor Killeen, Zeming Lin, Natalia Gimelshein, Luca Antiga, et al. Pytorch: An imperative style, high-performance deep learning library. *Advances in neural information processing systems*, 32, 2019.

[10] Hervé Turlier, Basile Audoly, Jacques Prost, and Jean-François Joanny. Furrow constriction in animal cell cytokinesis. *Biophysical journal*, 106(1):114–123, 2014.

[11] Stefan Van der Walt, Johannes L Schönberger, Juan Nunez-Iglesias, François Boulogne, Joshua D Warner, Neil Yager, Emmanuelle Gouillart, and Tony Yu. scikit-image: image processing in python. *PeerJ*, 2:e453, 2014.

[12] Pauli Virtanen, Ralf Gommers, Travis E Oliphant, Matt Haberland, Tyler Reddy, David Cournapeau, Evgeni Burovski, Pearu Peterson, Warren Weckesser, Jonathan Bright, et al. Scipy 1.0: fundamental algorithms for scientific computing in python. *Nature methods*, 17(3):261–272, 2020.
